# Comparative genomics reveals insight into the evolutionary origin of massively scrambled genomes

**DOI:** 10.1101/2022.05.09.490778

**Authors:** Yi Feng, Rafik Neme, Leslie Y. Beh, Xiao Chen, Jasper Braun, Michael Lu, Laura F. Landweber

**Author notes:** Author for Correspondence: Laura F. Landweber, Departments of Biochemistry and Molecular Biophysics and Biological Sciences, Columbia University, New York, NY, USA.

## Abstract

Ciliates are microbial eukaryotes that undergo extensive programmed genome rearrangement that converts long germline chromosomes into smaller gene-rich somatic chromosomes. Three well-studied ciliates include *Oxytricha trifallax*, *Tetrahymena thermophila* and *Paramecium tetraurelia*, but only the *Oxytricha* lineage has a massively scrambled genome whose assembly requires hundreds of thousands of precise DNA joining events. Here we study the emergence of genome complexity by examining the origin and evolution of discontinuous and scrambled genes in the *Oxytricha* lineage.

We sequenced, assembled and annotated the germline and somatic genomes of *Euplotes woodruffi* and the germline genome of *Tetmemena sp.*, and compared their genome rearrangement features to that of the model ciliate *Oxytricha trifallax*. The germline genome of *Tetmemena* is as massively scrambled and interrupted as *Oxytricha*’s: 13.6% of its gene loci rearrange via translocations and/or inversions. This study revealed that the earlier-diverged spirotrich, *E. woodruffi*, also has a scrambled genome, but approximately half as many loci (7.3%) are scrambled, supporting its position as a possible evolutionary intermediate in this lineage, in the process of accumulating complex genome rearrangements. Scrambled loci are more often associated with local duplications, supporting a simple model for the origin of scrambled genes via DNA duplication and decay.

## Introduction

Organisms do not always contain a single, static genome. Programmed genome rearrangements exist in many organisms, including ciliates (1), nematodes (2), lampreys (3) and zebra finches (4). Most cases involve excision, removal and rejoining of large regions of DNA to distinguish germline and somatic genomes. Ciliates are microbial eukaryotes with two types of nuclei: a somatic macronucleus (MAC) and a germline micronucleus (MIC). In the model ciliate *Oxytricha*, the MAC is entirely active chromatin (5), and the hub of transcription. The species in this study all have gene-sized “nanochromosomes” in the MAC, present at high copy number (6, 7, 8, 9, 10, 11). The diploid MIC, on the other hand, contributes to sexual reproduction, but its megabase-sized chromosomes are transcriptionally silent during asexual cell division.

Gene loci are often arranged discontinuously in the MIC, with short genic segments called Macronuclear Destined Sequences (MDSs), interrupted by stretches of non-coding DNA called Internally Eliminated Sequences (IESs) (Figure 1A). During sexual development, a new MAC genome rearranges from a copy of the zygotic MIC genome. MDSs join in the correct order and orientation, whereas MIC-limited genomic regions undergo programmed deletion, including repetitive elements, intergenic regions and IESs (Figure 1A). Though analogous to intron splicing, these events occur on DNA. The MDSs for some MAC chromosomes are called *scrambled* if they require translocation or inversion during MAC development (Figure 1A). Pairs of short repeats, called *pointers*, are present at MDS-IES junctions in both scrambled and nonscrambled loci (12, 13). Pointer sequences are present twice in the MIC, at the end of MDS *n* and the beginning of MDS *n+1*. One copy of the repeat is present at each MDS-MDS junction in a mature MAC chromosome (Figure 1A). These microhomologous regions help guide MDS recombination, but most are non-unique and the shortest pointers are just 2 bp. Thousands of long, noncoding template RNAs collectively program MDS joining (14, 15, 16).

**Figure 1.**
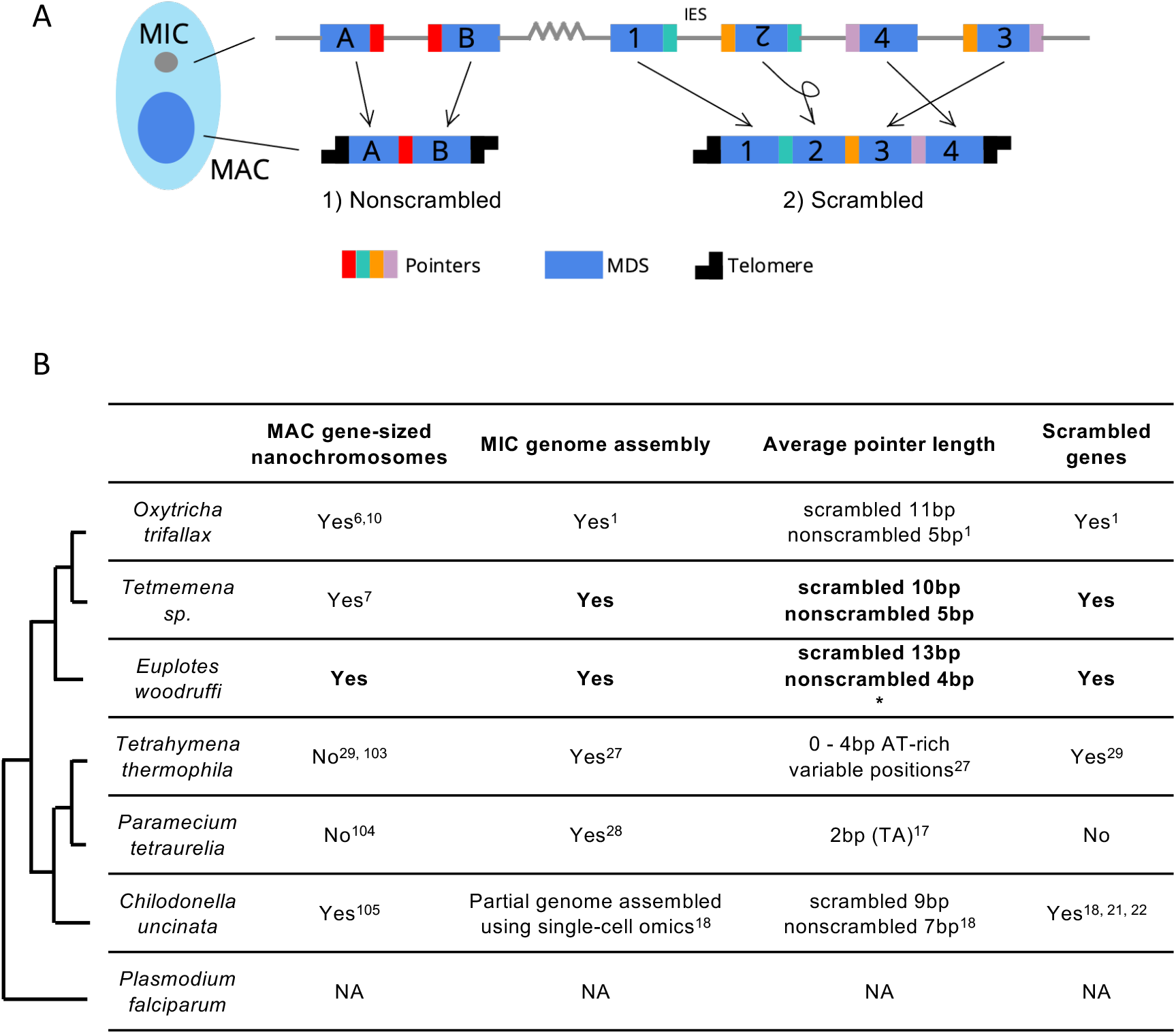
Genome rearrangements in representative ciliate species. A) Diagram of genome rearrangement in *Oxytricha*. Each ciliate cell contains a somatic macronucleus (MAC) and a germline micronucleus (MIC). During development, the MAC genome rearranges from a copy of the MIC genome. 1) Nonscrambled genes rearrange simply by joining consecutive macronuclear destined sequences (MDSs, blue boxes) and removing internal eliminated sequences (IESs, thin lines). 2) Rearrangement of scrambled genes requires MDS translocation and/or inversion. Pointers are microhomologous sequences (colored vertical bars) present in two copies in the MIC and only one copy in the MAC where consecutive MDSs recombine. B) Comparison of genome rearrangement features of representative ciliates and the non-ciliate *Plasmodium falciparum* as an outgroup (phylogeny adapted from (30, 31)). Conclusions from this study are shown in bold. * indicates that some scrambled pointers in *E. woodruffi* are much longer, as discussed in the results. Statistics for pointers <=30bp in *E. woodruffi* are shown.

Numerous studies have inferred the possible scope of genome rearrangement in different ciliate species using partial genome surveys. In *Paramecium*, PiggyMac-depleted cells fail to remove MIC-limited regions properly, which provided a resource to annotate ∼45,000 IESs prior to assembly of a complete draft MIC genome (17). The use of single-cell sequencing has allowed pilot studies to sample partial MIC genomes of diverse species (9, 18, 19, 20). Alignment of tentative MIC reads to either assembled MAC genomes or single-cell transcriptome data predicts over 20 candidate scrambled loci in two basal ciliates, *Loxodes sp*. and *Blepharisma americanum* (19) and hundreds of candidate loci in the tintinnid *Schmidingerella arcuata* (20). Nearly one third (31%) of approximately 5,000 surveyed transcripts may be scrambled in *Chilodonella uncinata* (18, Figure 1B), which has four confirmed cases of scrambled genes (21, 22). Transcriptome-based surveys offer less precise estimates, and cannot distinguish RNA splicing. Several computational pipelines have been developed to facilitate the inference of genome rearrangement features by split-read mapping in the absence of complete MIC or MAC reference genomes (23, 24, 25, 26). These useful tools provide partial insight to guide selection of species for full genome sequencing, which allows construction of complete rearrangement maps of a MIC genome onto a MAC genome for a reference species. Complete MIC genome reference assemblies are only currently available for three model ciliates: *Oxytricha trifallax* (1), *Tetrahymena thermophila* (27) and *Paramecium tetraurelia* (28).

*Oxytricha trifallax* has arguably the most exaggerated genome structure of any model organism (1), with programmed loss of over 90% of the germline MIC genome during development and massive descrambling to produce a MAC genome of over 18,000 chromosomes (10). This differs from the distantly-related *Tetrahymena* and *Paramecium* that both eliminate ∼30% of the germline genome (27, 28). *Paramecium* uses exclusively 2 bp pointers and lacks evidence of scrambled loci, which have been confirmed for four out of 2711 candidate scrambled loci in *Tetrahymena* (29, Figure 1B). *Tetrahymena and Paramecium* diverged from *Oxytricha* over one billion years ago (30, 31), which leaves a large gap in our understanding of the emergence of complex DNA rearrangements in the *Oxytricha* lineage.

Open questions include how did the *Oxytricha* germline genome acquire its high number of IES insertions and how do scrambled loci arise and evolve. Three previous studies tackled these questions at the level of single genes and orthologs, including DNA polymerase α, actin I and TEBPα (32, 33, 34, 35). Here, we provide the first comparative genomic analysis of *Oxytricha trifallax* and two other spirotrichous ciliates, *Tetmemena sp.* and *Euplotes woodruffi*. *Tetmemena sp.* is a hypotrich similar to *Tetmemena pustulata*, formerly *Stylonychia pustulata* (7), in the same family as *Oxytricha trifallax* (Figure 1B, 1, 7). Hypotrichs are noted for the presence of scrambled genes, based on previous ortholog comparisons (7, 32, 33, 34, 35, Figure 1B). *E. woodruffi,* together with the hypotrichous ciliates, belong to the class Spirotrichea (Figure 1B). Like hypotrichs, *Euplotes* also has gene-sized nanochromosomes in the MAC genome (8, 9, 36), but this outgroup uses a different genetic code (UGA is reassigned to cysteine, ref. 37) and little is known about its MIC genome. A partial MIC genome of *Euplotes vannus* was previously assembled, and it contains highly conserved TA pointers (9), consistent with previous observations in *Euplotes crassus* (38). This differs from *Oxytricha trifallax*, which uses longer pointers of varying lengths, with scrambled pointers typically longer than nonscrambled ones (1, Figure 1B). This observation suggests that longer pointers may supply more information to facilitate MDS descrambling, sometimes over great distances. Therefore, the preponderance of 2 bp pointers in the other *Euplotes* species could indicate limited capacity to support scrambled genes, and a partial genome survey of *E. vannus* concluded that at least 97% of loci are nonscrambled (9). Early studies of *Euplotes octocarinatus*, on the other hand, demonstrated its use of longer pointers (that usually contain TA) (39, 40), suggesting that some members of the *Euplotes* genus may have the capacity to support complex genome reorganization. To investigate the origin of scrambled genomes, we choose *E. woodruffi* as an outgroup, because it is closely related to *E. octocarinatus* (41) and feasible to culture in the lab.

We first present *de novo* assemblies of the micronuclear genome of *Tetmemena sp.* and both genomes of *E. woodruffi*. The availability of MIC and MAC genomes for both species allowed us to annotate and compare their genome rearrangement maps and other key features to each other and to *O. trifallax*. We find that transposable elements (TE) contribute substantially to differences in the MIC genomes sizes among these three species. The MIC genome of *Tetmemena* is extremely interrupted, like *Oxytricha*, but the *Euplotes* MIC genome is much more IES-sparse. We discovered the presence of 2429 scrambled genes in *E. woodruffi*, and present their gene architecture, with comparison to orthologous loci in the two hypotrich species and identify the presence of conserved pointers among orthologs for all three ciliates. We infer that the evolutionary origin of scrambled genes is associated with local duplications, providing strong support for a previously proposed simple evolutionary model requiring only duplication and decay (42). We also identify examples of non-coding regions that can be eliminated as either IESs or introns, suggesting the opportunity to delete some sequences at either the DNA or RNA level (33). Finally, we investigated the evolution of some of the most complex nested loci, with segments for multiple genes embedded between those for other genes, so-called Russian doll loci. We find a high level of synteny between Russian doll genes in *Oxytricha* and *Tetmemena*, with species-specific transposon insertions contributing to complex germline architecture.

## Results

### Germline genome expansion via repetitive elements

*Tetmemena sp.* and *E. woodruffi* were both propagated in laboratory culture from single cells. The *E. woodruffi* MAC genome was sequenced and assembled from paired-end Illumina reads from whole cell DNA, which is mostly MAC-derived. For comparative analysis, the MAC genome of *E. woodruffi* was assembled using the same pipeline previously used for *Tetmemena sp.* (7). Because MIC DNA is significantly more sparse than MAC DNA in individual cells (13) MIC DNA was enriched before sequencing (see Methods); however, this leads to much lower sequence coverage of the MIC than the MAC. Third-generation long reads (Pacific Biosciences and Oxford Nanopore Technologies) were combined with Illumina paired-end reads (Methods, see genome coverage in Table S1) to construct hybrid genome assemblies for *Tetmemena sp.* and *E. woodruffi*. Though the final genome assemblies are still fragmented, often due to transposon or other repetitive insertions at boundaries (Figure S1), the current draft assemblies cover most (>90%) MDSs for 89.1% of MAC nanochromosomes in *Tetmemena*, and for 90.0% of MAC nanochromosomes in *E. woodruffi*. This allowed us to establish near-complete rearrangement maps for the newly assembled genomes of *Tetmemena* and *E. woodruffi,* at a level comparable to the published reference for *O. trifallax* (1), which is appropriate for comparative analysis.

Table 1 shows a comparison of genome features for the three species. The three MAC genomes are similar in size, with most nanochromosomes encoding only one gene. Like *O. trifallax* (6), the maximum number of genes encoded on one chromosome is 7-8 (Table 1). Surprisingly, the MIC genome sizes differ substantially: the *Tetmemena* MIC genome assembly is 237 Mbp, nearly half that of *Oxytricha*. The *E. woodruffi* MIC genome assembly is even smaller, approximately 172 Mbp (Table 1).

**Table 1.** Statistics of MAC and MIC genomes in three species

|  | *Oxytricha trifallax* |  | *Tetmemena sp.* |  | *Euplotes woodruffi* |  |
| --- | --- | --- | --- | --- | --- | --- |
|  | MAC<sup>6,b</sup> | MIC<sup>1</sup> | MAC<sup>7</sup> | MIC | MAC | MIC |
| genome size (Mbp) | 67.1 | 496 | 60.6 | 237 | 72.2 | 172 |
| N50 (bp) | 3,745 | 27,807 | 3,339 | 14,722 | 2,702 | 44,656 |
| GC% | 31.36 | 28.44 | 37.05 | 32.17 | 36.56 | 35.31 |
| number of contigs<sup>a</sup> | 22,426 | 25,720 | 25,206 | 28,446 | 35,099 | 17,655 |
| Two-telomere contigs | 14,225 | - | 15,802 | - | 19,061 | - |
| Telomeric contigs | 20,336 | - | 21,165 | - | 28,294 | - |
| Single-gene telomeric contigs | 76.1% | - | 75.5% | - | 68.5% | - |
| Maximum number of genes on a telomeric contig | 8 | - | 7 | - | 8 | - |

<sup>a</sup> TBE contaminants in MAC contigs were removed (Methods). Therefore, 24 *Oxytricha* MAC contigs and 13 *Tetmemena* MAC contigs were removed from the published versions.
<sup>b</sup> This study used the MAC genome of *Oxytricha* from Swart et al. (6) instead of the long-read assembly in Lindblad et al. (10), because the short MAC genomes in the present study were primarily assembled from Illumina reads, as in (6). (10) updated (6) by including nanochromosomes captured in single long reads, which are currently not available for the other two species. The MIC genomes of *Tetmemena* and *E. woodruffi* were assembled to a similar N50 as the reference *O. trifallax* genome (1) for comparative analysis.

The expansion of repetitive elements in the *Oxytricha* lineage may contribute to the difference in MIC genomes sizes (Figure 2A-C). *Oxytricha* has a variety of TEs in the MIC, with telomere-bearing elements (TBEs) of the Tc1/*mariner* family the most abundant (1, 43, Table S2). A complete TBE transposon contains three open reading frames (ORFs). ORF1 encodes a 42kD transposase with a DDE-catalytic motif. Though present only in the germline, TBEs are so abundant in hypotrichs that some were partially recovered and assembled from whole cell DNA (42). The *Oxytricha* MIC genome contains ∼10,000 complete TBEs and ∼24,000 partial TBEs, which occupy approximately 15.20% (75 Mbp) of the genome (Figure 2A, Table S3, 1, 42). *Tetmemena*, on the other hand, has many fewer TBE ORFs and only 48 complete TBEs (Table S3), comprising 1.83% (4.3 Mbp) of its MIC genome (Figure 2B). *Euplotes crassus* has also been reported to have an abundant transposon family called Tec elements (<u>T</u>ransposon of *<u>E</u>uplotes <u>c</u>rassus*). Like TBEs, each Tec consists of three ORFs, and ORF1 also encodes a transposase from the Tc1/*mariner* family (44, 45, 46, 47, 48). Using the three ORFs of Tec1 and Tec2 as queries for search, we identified 74 complete Tec elements in *E. woodruffi*. Collectively, Tec ORFs occupy 3.6 Mbp, corresponding to only 2.1% of the MIC genome (Figure 2C). Notably, the transposase-encoding ORF1, previously implicated in genome rearrangement in *Oxytricha* (49) is more abundant than the other two TBE/Tec ORFs in all three ciliates (Table S3).

**Figure 2.**
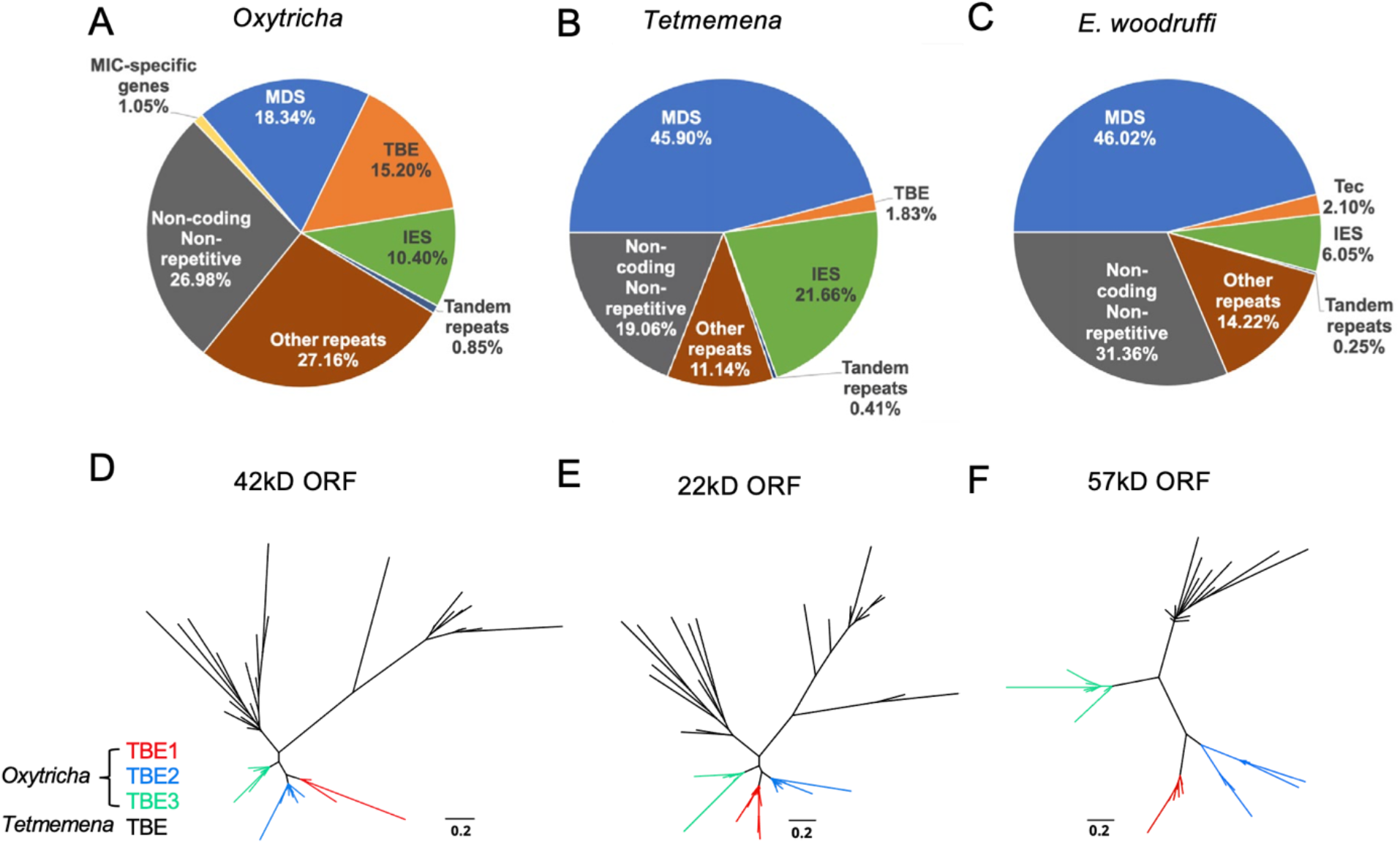
The three MIC genomes differ in repeat content, especially transposable elements. A-C) MIC genome categories for (A) *Oxytricha trifallax*, (B) *Tetmemena sp.*, and (C) *E. woodruffi*. *Oxytricha* displays the greatest proportion of repetitive elements (TBE, Other repeats, and Tandem repeats) relative to the other species. *Oxytricha* MIC-specific genes were annotated in (1, 102). D-F) Phylogenetic analysis of the three TBE ORFs in *Oxytricha* and *Tetmemena*: (D) 42kD, (E) 22kD, and (F) 57kD, suggest that TBE3 (green) is the ancestral transposon family in *Oxytricha*. For each ORF, 30 protein sequences from each species were randomly subsampled and Maximum Likelihood trees constructed using PhyML (97).

*Oxytricha* contains three families of TBEs. TBE3 appears to be the most ancient among hypotrichs, based on previous analysis of limited MIC genome data (43). We constructed phylogenetic trees using randomly subsampled TBE sequences for all three ORFs from *Oxytricha* and *Tetmemena* (Figure 2D-F). This confirmed that only TBE3 is present in the *Tetmemena* MIC genome, as proposed in (43). This also suggests that TBE1 and TBE2 expanded in *Oxytricha* after its divergence from other hypotrichous ciliates. As illustrated in Figure S1, the MIC genome contexts of TBEs in *Oxytricha* and *Tetmemena* are similar, with many TE insertions within IESs, suggesting that some TE insertions may have generated IESs, as demonstrated in *Paramecium* (50, 51). For further discussion of the conservation of TBE locations, see the section, “*Oxytricha* and *Tetmemena* share conserved rearrangement junctions’’ below.

Additionally, Repeatmodeler/Repeatmasker identified that *Oxytricha* has more MIC repeats in the “Other” category than *Tetmemena* or *E. woodruffi* (Figure 2, subcategories of repeat content in Table S2). 214 Mbp of the *Oxytricha* MIC genome (43%, which is greater than 35.9% reported in ref. 1 that used earlier versions of the software) is considered repetitive (including TBEs, tandem repeats and other repeats in Figure 2), versus 31.7 Mbp for *Tetmemena* (13.4%) and 28.5 Mbp (16.8%) for *E. woodruffi*. *Oxytricha*’s additional ∼180 Mbp in repeat content partially explains the significantly larger MIC genome size of *Oxytricha* versus the other spirotrich ciliates.

### The *E. woodruffi* genome has fewer IESs

We used the genome rearrangement annotation tool SDRAP (52) to annotate the MIC genomes of *Oxytricha, Tetmemena* and *E. woodruffi* (Methods). Consistent with their close genetic distance, the genomes of *O. trifallax* and *Tetmemena* have similarly high levels of discontinuity (Figure 3A): We annotated over 215,299 MDSs in *Oxytricha* and over 215,624 in *Tetmemena* MDSs with similar MDS length distributions (Figure 3A). By contrast, *E. woodruffi* MDSs are typically longer, which indicates a less interrupted genome (Figure 3A). We compared the number of MDSs between single-copy orthologs for single-gene MAC chromosomes across the three species and found that the orthologs have similar lengths (Figure S2). There is a strong positive correlation between number of MDSs for orthologous genes in *Oxytricha* and *Tetmemena* (R^2^ = 0.75, Figure 3B). There is no correlation among number of MDSs between orthologs of *E. woodruffi* and *Oxytricha* (R^2^ = 0.003, Figure 3C), since *E. woodruffi* orthologs typically contain fewer MDSs.

**Figure 3.**
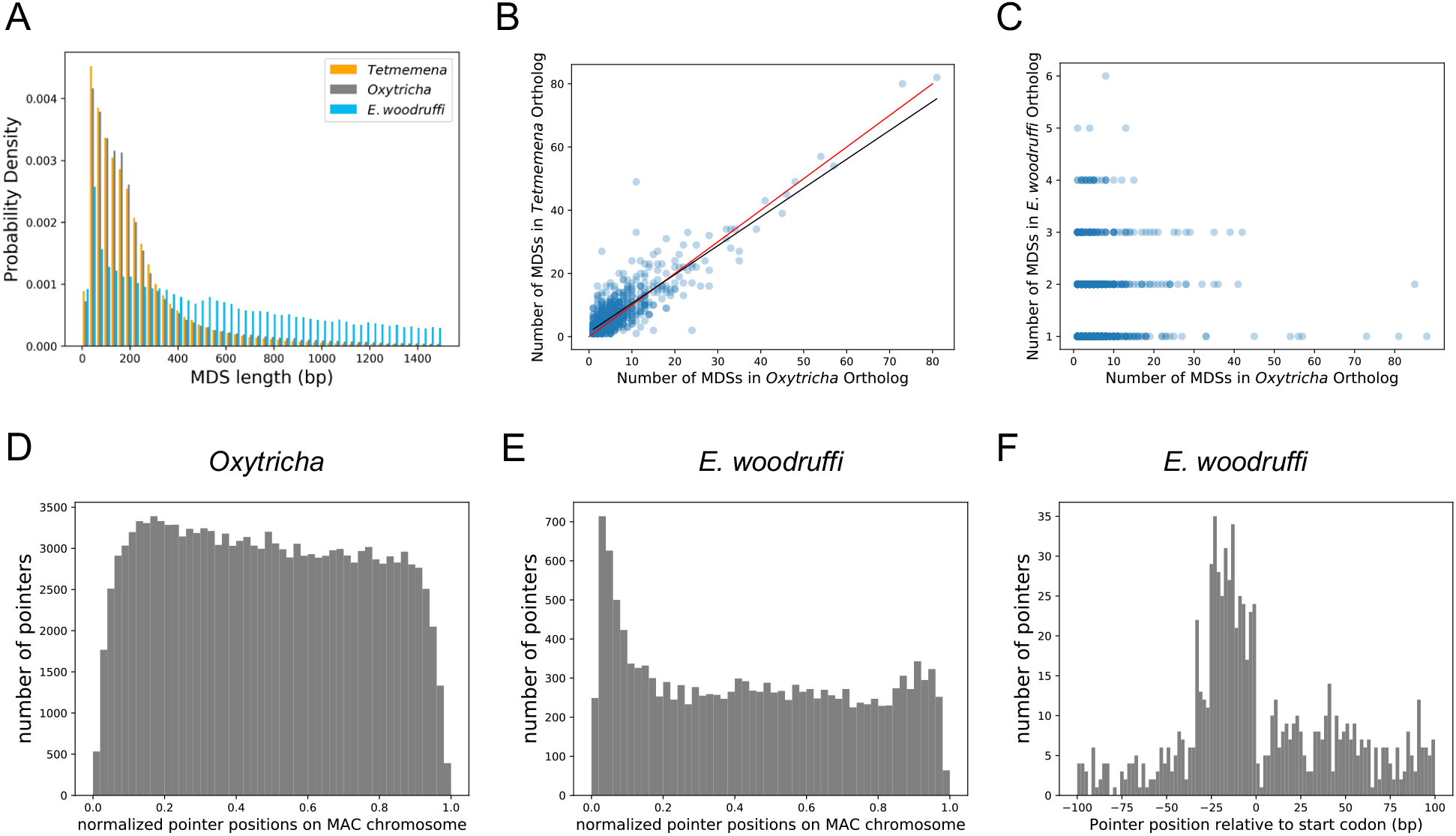
The three MIC genomes are interrupted by IESs at different levels. A) MDSs of *E. woodruffi* are longer compared to *Oxytricha* or *Tetmemena*. B) Positive correlation between the numbers of MDSs for orthologous genes in *Tetmemena* and in *Oxytricha* for 903 single-gene orthologs. Black line is the function of linear regression (R^2^ = 0.75). Red line is y=x. C) Orthologs in *E. woodruffi* have fewer MDSs compared to *Oxytricha*, with no correlation (R^2^ = 0.003). Note that many highly discontinuous genes in *Oxytricha* are IES-less in *E. woodruffi* (present on one MDS). 917 single-gene orthologs are shown. D) Distribution of pointers on single-gene MAC chromosomes in *Oxytricha vs.* E) *E. woodruffi*, with MAC chromosomes oriented in gene direction. Pointers significantly accumulate at the 5’ end of single-gene MAC chromosomes in *E. woodruffi*. (F) Pointer positions on 3684 two-MDS MAC chromosomes demonstrate a preference upstream of the start codon.

The *E. woodruffi* genome is generally much less interrupted than that of *Oxytricha* or *Tetmemena*. 39.9% of MAC nanochromosomes in *E. woodruffi* lack IESs (IES-less nanochromsomes) compared to only 4.1% and 4.4% in *Oxytricha* and *Tetmemena*, respectively. The sparse IES distribution (as measured by plotting pointer distributions) in *E. woodruffi* displays a curious 5’ end bias on single-gene MAC chromosomes, oriented in gene direction (Figure 3E). A weak 5’ bias is also present in *Oxytricha* (Figure 3D) and *Tetmemena* (not shown). In addition, *E. woodruffi* IESs preferentially accumulate in the 5’ UTR, a short distance upstream of start codons (Figure 3F). Notably, the median distance between the 5’ telomere and start codon in *E. woodruffi* is just 54 bp for single-gene chromosomes.

### *E. woodruffi* has an intermediate level of genome scrambling

Scrambled genome rearrangements exist in all three species, which we report here for the first time in *Tetmemena* and the early-diverged *E. woodruffi*. Previous studies have described scrambled genes with confirmed MIC-MAC rearrangement maps for a limited species of hypotrichs (1, 7, 32, 33, 34, 35) and *Chilodonella* (21, 22), but not in *Euplotes*. Consistent with the phylogenetic placement of *Euplotes* as an earlier-diverged outgroup to hypotrichs (53, 54), the *E. woodruffi* genome is scrambled, but it contains approximately half as many scrambled genes (2429 genes encoded on 1913 chromosomes, or 7.3% of genes), versus 15.6% scrambled in *Oxytricha trifallax* (3613 genes encoded on 2852 chromosomes) and 13.6% in *Tetmemena* (3371 genes encoded on 2556 chromosomes). The *E. woodruffi* lineage may therefore reflect an evolutionary intermediate stage between ancestral genomes with only modest levels of genome scrambling versus the more massively scrambled genomes of hypotrichs.

We infer that many genes were likely scrambled in the last common ancestor of *Oxytricha* and *Tetmemena*, because these two species share approximately half of their scrambled genes (Table S4). The majority of *Oxytricha/Tetmemena* scrambled genes have no detectable ortholog in *E. woodruffi*, and approximately 80% of *E. woodruffi* scrambled genes have no ortholog in the other two species. This could be explained by a scenario in which either the property of gene scrambling contributes to *de nov*o gene birth, or nucleotide divergence obscures their orthology. The latter is more plausible because most nonscrambled genes in *Oxytricha/Tetmemena* also have no detectable ortholog in *E. woodruffi* (Table S4). Furthermore, the majority of these scrambled genes are not new genes, since they possess at least one ortholog in other ciliate species (Table S4).

### Scrambled genes are associated with local paralogy

Notably, scrambled genes in all three species generally have more paralogs (Figure 4). We identified orthogroups containing genes derived from the same gene in the last common ancestor of the three species (Methods). For each species, orthogroups with at least one scrambled gene are significantly larger than those containing no scrambled genes (*p*-value <1e-5, Mann-Whitney U test, Figure 4A-C). This association suggests a possible role of gene duplication in the origin of scrambled genes.

**Figure 4.**
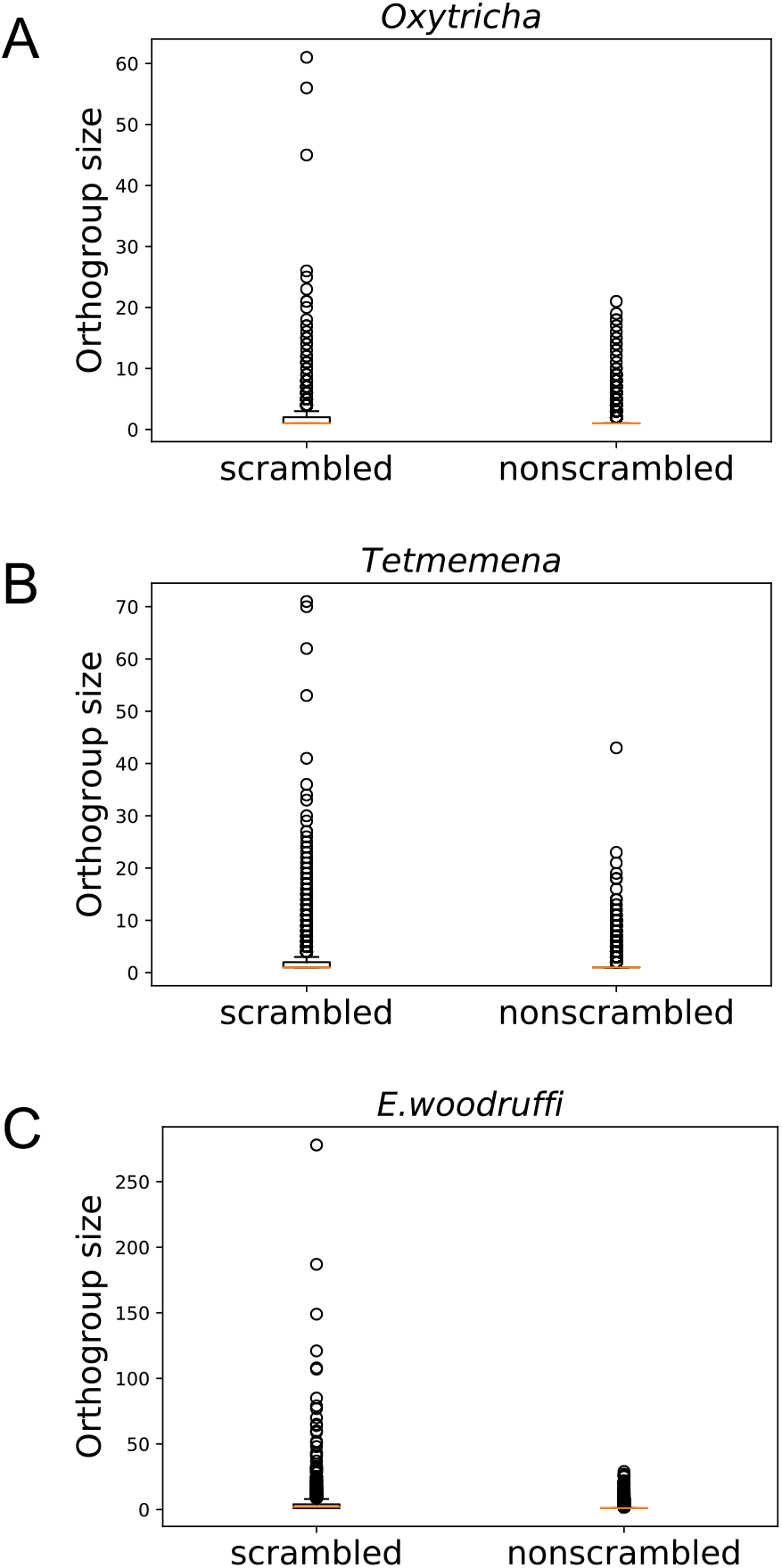
Scrambled genes have more paralogs than nonscrambled genes in the three species. Orthogroups containing at least one scrambled gene (“scrambled”) are larger than orthogroups that lack scrambled genes (“nonscrambled”) in A) *Oxytricha*, B) *Tetmemena* and C) *E. woodruffi*.

Scrambled pointers are generally longer than nonscrambled ones in all three species (Figure S3), consistent with prior observations (1) and the possibility that longer pointers participate in more complex rearrangements, including recombination between MDSs separated by greater distances (55). Scrambled and nonscrambled IESs also differ in their length distribution (Figure S3). Notably, scrambled “pointers” in *E. woodruffi* can be as long as several hundred base pairs (median 48 bp, average 212 bp) unlike the more typical 2-20 bp canonical pointers. These long “pointers” in *E. woodruffi* are more likely partial MDS duplications (Figure S4A). We also identified MDSs that map to two or more paralogous regions within the same MIC contig (Table S5), therefore representing MDS duplications and not alleles. Such paralogous regions could be alternatively incorporated into the rearranged MAC product. Notably, we find that, for all three species, there are significantly more scrambled than nonscrambled MAC chromosomes that contain at least one paralogous MDS (chi-square test, *p*- value <1e-10; Table S5). An example is shown in Figure S4A (MDS 7 and 7’).

The presence of paralogous MDSs can contribute to the origin of scrambled rearrangements, as proposed in an elegant model by Gao et al. (ref.42; illustrated in Figure S4B). The model proposes that initial MDS duplications permit alternative use of either MDS copy into the mature MAC chromosome. As mutations accumulate in redundant paralogs, cells that incorporate the least decayed MDS regions into the MAC gene would have both a fitness advantage and a better match to the template RNA (14) that guides rearrangement, thus increasing the likelihood of incorporation into the MAC chromosome. The paralogous regions containing more mutations would gradually decay into IESs and scrambled pointers eventually reduced to a shorter length. The extended length “pointers” that we identified in *E. woodruffi* may reflect an intermediate stage in the origin of scrambled genes (Figure S4B).

This model may generally explain the abundance of “odd-even” patterns in ciliate scrambled genes (55, 56). As illustrated in Figure S4A, the even- and odd-numbered MDSs for many scrambled genes derive from different MIC genome clusters. The model predicts that the IES between MDS *n-1* and *n+1* often derives from ancestral duplication of a region containing MDS *n* (Figure S5A). To test this hypothesis explicitly, we extracted from all odd-even scrambled loci in the three species all sets of corresponding MDS/IES pairs that are flanked by identical pointers on both sides, i.e. all pairs of scrambled MDSs and IESs, where the IES between MDS *n-1* and *n+1* is directly exchanged for MDS *n* during DNA rearrangement (S1 and S2 in Figure S5A). To exclude the possibility of alleles confounding this analysis, MDS and IES pairs were only considered if they map to the same MIC contig. In *E. woodruffi*, the lengths of these MDS/IES pairs strongly correlate (Spearman correlation ρ=0.755, *p*<1e-5, Figure S5B). Moreover, many MDS and IES sequence pairs also share sequence similarity, consistent with paralogy: For 248 MDS-IES pairs of similar length, 90.3% share a core sequence with ∼97.5% identity across 8-100% of both the IES and MDS length. The lowest end of these observations is also compatible with an alternative model (33) in which direct recombination between IESs and MDSs at short repeats can lead to expansion of odd-even patterns. For *Oxytricha* and *Tetmemena*, the MDS and IES lengths for such MDS/IES pairs also correlate positively, but more moderately (Figure S5D-E). Remarkably, the odd-even-containing loci that are species-specific, and therefore became scrambled more recently, have the strongest length correlation (Figure S5C-E) and more pairs that display sequence similarity (Table S6) relative to older loci (scrambled in two or more species). This result is consistent with an evolutionary process in which mutations accumulate in one copy of the MDS, gradually obscuring its sequence homology and ability to be incorporated as a functional MDS, and eventually its ability to be recognized by the template RNAs that guide DNA rearrangement. This analysis also suggests that most of the odd-even scrambled loci in *E. woodruffi* arose recently, because there is greater sequence similarity between MDSs and the corresponding IESs that they replace. Conversely, we infer that most loci that are scrambled in both *Oxytricha* and *Tetmemena* became scrambled earlier in evolution, since they display weaker sequence similarity between exchanged MDS and IES regions.

### *Oxytricha* and *Tetmemena* share conserved DNA rearrangement junctions

To understand the conservation of genome rearrangement patterns, we developed a pipeline guided by protein sequence alignment to compare pointer positions for orthologous genes between any two species (Methods, Figure 5A). We compared pointers for 2503 three-species single-copy orthologs. 4448 pointer locations are conserved between *Oxytricha* and *Tetmemena* on 1345 ortholog pairs (Table S7), representing 38.3% of pointers in these orthologs in *Oxytricha* and 30.9% in *Tetmemena*. For *Oxytricha*/*E. woodruffi* and *Tetmemena*/*E. woodruffi* comparisons, 56 and 58 pointer pairs are conserved, respectively. We also identified 23 pointer locations shared among all three species (Table S7, Figure 5B, Data S1).

**Figure 5.**
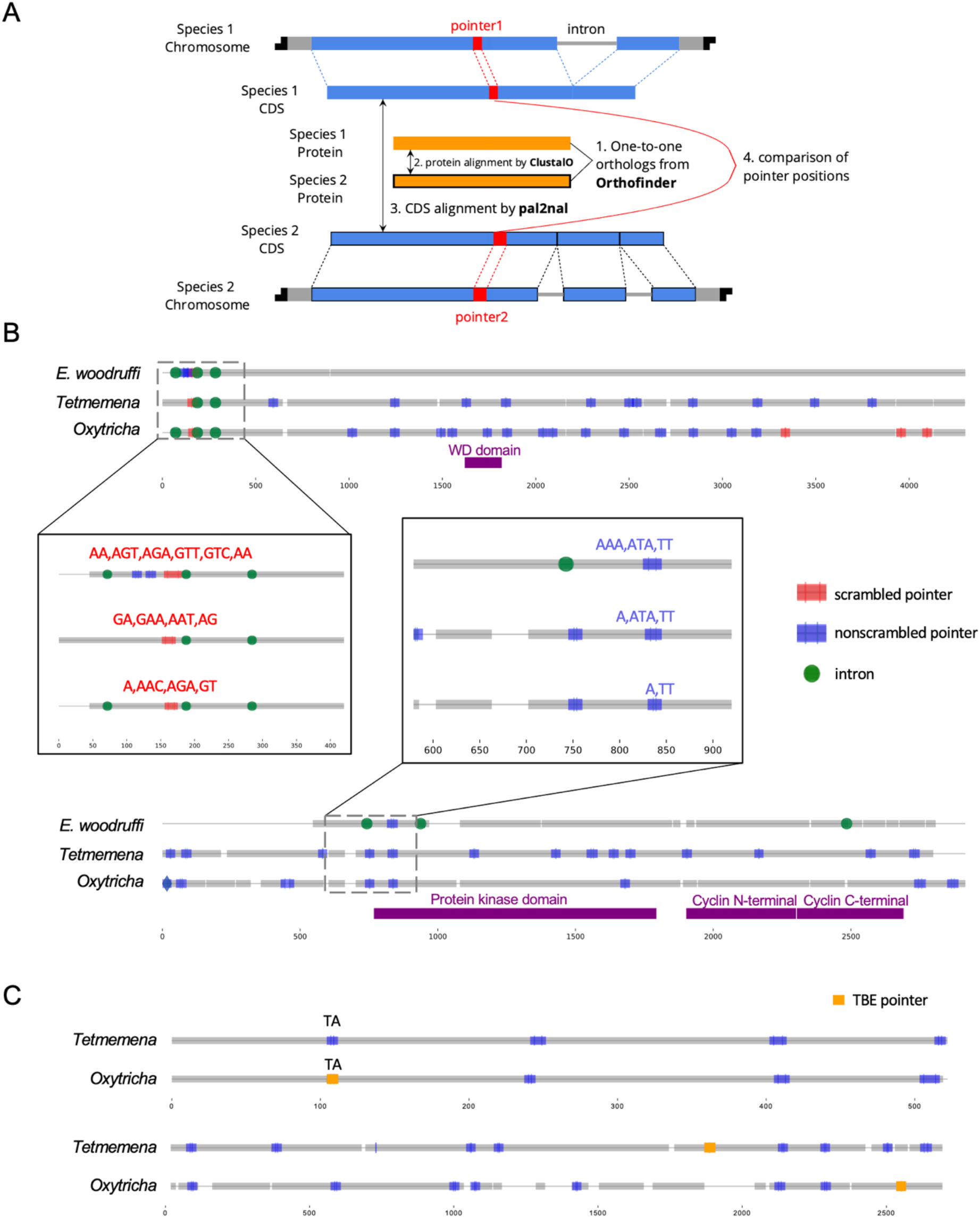
Identification and examples of conserved pointers. A) Pipeline for comparison of pointer positions in orthologs. Orthologs are first grouped by OrthoFinder (98) and protein sequences of single-copy orthologs aligned by Clustal Omega (99). Then the protein alignments are reverse translated to coding sequence (CDS) alignments by a modified script of pal2nal (100, Methods). Pointers are annotated on the CDS alignments for comparison between any two orthologs. B) Two examples of pointer conservation across three species. Gray lines represent the alignment of orthologous CDSs and boxes show magnified regions containing conserved pointers. The top panel shows a conserved scrambled pointer (*Oxytricha*: Contig889.1.g68; *Tetmemena*: LASU02015390.1.g1; *E. woodruffi*: EUPWOO_MAC_30105.g1). The bottom panel shows a conserved nonscrambled pointer (*Oxytricha*: Contig19750.0.g98; *Tetmemena*: LASU02002033.1.g1; *E. woodruffi*: EUPWOO_MAC_31621.g1). Pointer sequences are noted and commas indicate reading frame. Protein domains detected by HMMER (101) are marked in purple. C) Examples of TBE insertions in nonscrambled IESs. The upper pair of sequences show an *Oxytricha* TBE pointer (orange insertion of an incomplete TBE2 transposon containing the 42kD and 57kD ORFs) conserved with a *Tetmemena* non-TBE pointer (*Oxytricha*: Contig736.1.g130; *Tetmemena*: LASU02012221.1.g1). Both species have a TA pointer at this junction. The bottom pair of sequences illustrate a case of nonconserved TBE pointers (*Oxytricha*: Contig17579.0.g71; *Tetmemena*: LASU02007616.1.g1).

To test if these pointer locations are genuinely conserved versus coincidental matching by chance, we performed a Monte Carlo simulation, as also used to study intron conservation (57). We randomly shuffled pointer positions on CDSs 1000 times, and counted the number of conserved pointer pairs expected for each simulation (Methods). Of the 1000 simulations, none exceeded the observed number of conserved pointer pairs between *Oxytricha* and *Tetmemena* (*p*-value < 0.001), suggesting evolutionary conservation of pointer positions (Table S7). A similar result was obtained for pointers conserved in all three species (Table S7). However, the numbers of pointer pairs conserved between *Oxytricha*/*E. woodruffi* and *Tetmemena*/*E. woodruffi* are similar to the expectations by chance (Table S7). The low level of pointer conservation of either hypotrich with *E. woodruffi* may reflect the smaller number of IESs in *E. woodruffi*; hence most pointers would have arisen in the hypotrich lineage. Furthermore, *E. woodruffi* is genetically more distant from the two hypotrichs; hence the accumulation of substitutions would obscure protein sequence homology, which we used to compare pointer locations. For ortholog pairs between *Oxytricha* and *Tetmemena*, scrambled pointers are significantly more conserved than nonscrambled ones (chi-square test, *p*-value <1e-10, Table S8). We also find that most pointer sequences differ even if the positions are conserved (Figure 5B, Data S1, Table S9), suggesting that substitutions may accumulate in pointers without substantially altering rearrangement boundaries.

*Oxytricha* and *Tetmemena* both contain a high copy number of TBE transposons (1, 43, Table S3). We investigated the level of TBE conservation between these two species. To identify orthologous insertions, we focus on TBE insertions in nonscrambled IESs on single-copy orthologs, which include 1706 *Oxytricha* TBEs inserted in 1296 nonscrambled IESs (multiple TBEs can be inserted into an IES) and 180 *Tetmemena* TBEs inserted into 170 nonscrambled IESs. We refer to the pointer flanking a TBE-containing IES as a *TBE pointer*. No TBE pointer locations are conserved between two species. This suggests that TBEs might invade the genomes of *Oxytricha* and *Tetmemena* independently, or still be actively mobile in the genome. Only 27 *Oxytricha* TBE pointers (containing 36 TBEs) are conserved with non-TBE pointers in *Tetmemena* (Data S2, Figure 5C). No *Tetmemena* TBE pointer is conserved with an *Oxytricha* non-TBE pointer. This suggests that TBE insertions may preferentially produce new rearrangement junctions instead of inserting into an existing IES.

### Intron locations sometimes coincide with DNA rearrangement junctions

Among three-species orthologs, intron locations sometimes map near pointer positions (Figure 5B, Figure S6). IESs and introns are both non-coding sequences removed from mature transcripts, though at different stages. A previous single-gene study observed that an IES in *Paraurostyla* overlaps the position of an intron in *Uroleptus*, *Urostyla* and also the human homolog (33). This observation suggested an intron-IES conversion model in which the ability to eliminate non-coding sequences as either DNA or RNA provides a potential backup mechanism. Such interconversion has also been observed between two strains of *Stylonychia* (58). In the present study, we identified 174 potential examples of intron-IES conversion in the three species (Figure S6, Table S10): 103 (59.2%) *E. woodruffi* introns map near *Oxytricha*/*Tetmemena* pointers. A Monte Carlo simulation for these intron-IES comparisons (Table S10) revealed that *p* <0.001 for most three-species comparisons. For two-species comparisons, only the comparison between *Oxytricha* intron positions vs. *Tetmemena* IES junctions was significant (*p =* 0.008) (Table S11). Notably, *Tetmemena* intron locations rarely coincide with *Oxytricha* IESs (Table S11), suggesting a possible bias in the direction of intron-IES conversion during evolution.

### Evolution of complex genome rearrangements: Russian doll genes

Genome rearrangements in the *Oxytricha* lineage can include overlapping and nested loci, with MDSs for different MAC loci embedded in each other (1, 59). When multiple gene loci are nested in each other, these have been called Russian doll loci (59). *Oxytricha* contains two loci with five or more layers of nested genes (59). *Oxytricha* and *Tetmemena* display a high degree of synteny and conservation in both Russian doll loci. In the first Russian doll gene cluster, one nested gene (green) is present in *Oxytricha* but absent in *Tetmemena* (Figure 6A, Figure S7, Figure S8), confirmed by PCR (Methods). *Oxytricha* also has a complete TBE3 insertion in the green gene (Figure 6A, Figure S7A), hinting at a possible link between transposon and new gene insertion. In addition, a two-gene chromosome in *Oxytricha* (orange) is present as two single-gene chromosomes in *Tetmemena* (Figure 6A, Figure S7). In *Oxytricha*, seven orange MDSs ligate across two loci via an 18 bp pointer (TATATCTATACTAAACTT) to form a 2-gene nanochromosome. However, in *Tetmemena*, telomeres are added to the ends of both gene loci instead, forming two independent MAC chromosomes (Figure 6A, Figure S7). The second Russian doll locus has an example of a long, conserved pointer (orange dotted line) that bridges three other loci (the green and blue scrambled loci and one nonscrambled locus, Figure 6B). Close to this region is a decayed TBE insertion (769bp) in *Oxytricha.* None of the *E. woodruffi* orthologs of both Russian doll loci map to the same MIC contig, which suggests that the Russian doll cluster arose after the divergence of *Euplotes* from the common ancestor of *Oxytricha* and *Tetmemena*.

**Figure 6.**
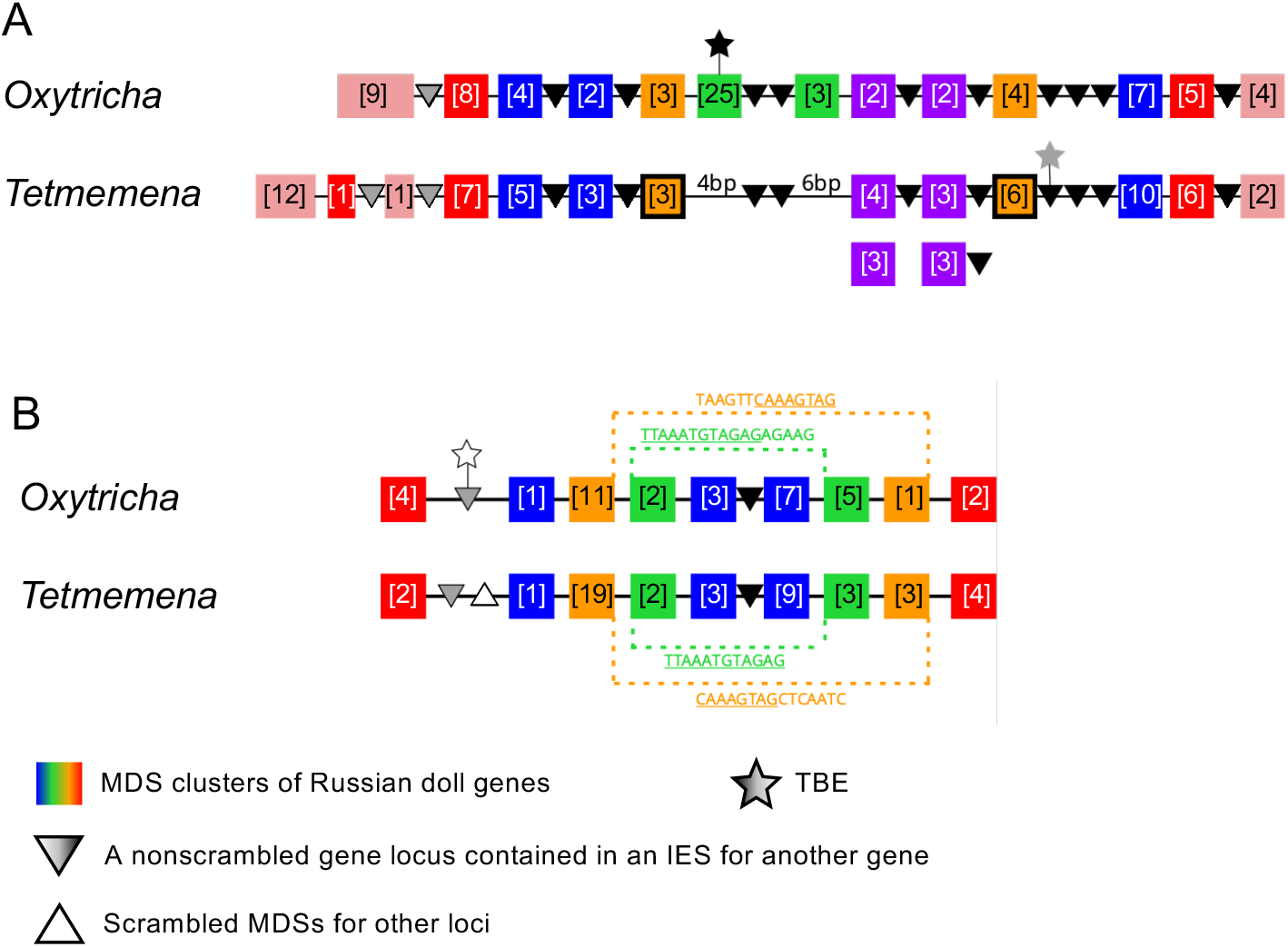
Synteny in “Russian doll” loci in *Oxytricha* and *Tetmemena*. A) Schematic comparison of the Russian doll gene cluster on *Oxytricha* MIC contig OXYTRI_MIC_87484 vs. *Tetmemena* MIC contig TMEMEN_MIC_21461. Boxes of the same color represent clusters of MDSs for orthologous genes (detailed map in Figure S7 and Figure S8). Numbers in brackets indicate the number of MDSs in each cluster, grouped by MAC chromosome. One nested gene (green) in *Oxytricha* is absent from *Tetmemena*. A two-gene chromosome (orange) that derives from seven MDSs in *Oxytricha* is processed as two single-gene chromosomes in *Tetmemena* instead (indicated by black border around orange boxes). The purple gene in *Oxytricha* has two paralogs in *Tetmemena*. Black triangles represent conserved, orthologous, nonscrambled gene loci inserted between nested Russian doll genes. Empty triangle represents scrambled MDSs for other loci. Gray triangles, complete nonscrambled MAC loci embedded between gene layers in one species with no orthologous gene detected in the other species. Black star, a complete TBE transposon insertion. Gray star, a partial TBE insertion. B) *Oxytricha* MIC contig OXYTRI_MIC_69233 vs. *Tetmemena* MIC contig TMEMEN_MIC_22886. Pointer sequences bridging the nested MDSs of orange and green genes are highlighted. The underlined pointer portions are conserved between species, e.g. the last 8bp of the *Oxytricha* pointer, <u>T</u>AAGTTCAAAGTAG, are identical to the first 8bp of <u>C</u>AAAGTAGCTCAATC in *Tetmemena*, illustrating pointer sliding (35), or gradual shifting of MDS/IES boundaries. White star indicates a decayed TBE with no ORF identified.

## Discussion

The highly diverse ciliate clade provides a valuable resource for evolutionary studies of genome rearrangement. However, complete assembly and annotation of germline MIC genomes has concentrated on the model ciliates *Tetrahymena, Paramecium* and *Oxytricha*. To provide insight into genome evolution in this lineage, we assembled and compared the complete germline and somatic genomes of *Tetmemena sp*. and an outgroup, *Euplotes woodruffi*, to that of *Oxytricha trifallax*. This expands our knowledge of the diversity of ciliate genome structures and the evolutionary origin of complex genome rearrangements.

Dramatic variation in transposon copy number (TBE and Tec elements) from the Tc1/*mariner* family appears to explain most of the variation in MIC genome size. In many eukaryotic taxa, genome size can differ dramatically even for closely related species, a phenomenon known as the “C-value paradox” (60). Our present observations are compatible with previous reports that the repeat content of the genome, especially transposon content, positively correlates with genome size (61).

*Oxytricha* has three TBE families in the MIC genome, but only TBE3 is present in *Tetmemena*. This agrees with our previous inference that TBE3 is present at the base of the transposon lineage in hypotrichous ciliates, representing the ancestral state (43). Tens of thousands of TBE1 and TBE2 transposons then expanded specifically in *Oxytricha*, due to either a more recent invasion or, alternatively, mutation of a TBE3 in the *Oxytricha* lineage to produce its successful TBE 1 and 2 elements. RNAi silencing of all three, but not individual, TBE transposase types leads to stalled genome rearrangement in *Oxytricha*, which suggests involvement of the encoded transposase in genome differentiation (49). This also suggests the possibility that some TBEs could have specialized roles. Despite a high copy number of TBEs in both *Oxytricha* and *Tetmemena*, no TBE is identical among TE insertions in nonscrambled IESs in our analysis. Even the two Russian doll regions with high synteny have non-conserved TBEs and TBE insertions appear together with insertion or deletion of nested genes. These observations suggest that TBEs may be active in these genomes and contribute to the evolution of genome structure.

Curiously, we observed a higher copy number of the transposase-encoding ORF1 in TBE and Tec elements compared to the other two ORFs for all three genomes (Table S3). This observation raises the possibility that the transposase may function independently of the other two downstream genes, consistent with our previous conclusions that the TBE transposase may participate in genome rearrangement in *Oxytricha* (49), possibly contributing to DNA cleavage. However, it is not clear if and how the other two TBE-encoded genes may be involved. The 22kD ORF2 has unknown function and the 57kD ORF3 contains zinc finger and kinase domains (43). In Tec elements, the ∼57kD ORF2 encodes a tyrosine-type recombinase (62) and the 20kD ORF3 has unknown function (46). All three *Oxytricha* TBE ORFs are under purifying selection, suggesting function (43).

In the relatively IES-poor genome of *E. woodruffi*, IESs accumulate upstream of start codons, similar to the 5’ bias of introns in intron-poor organisms (63). The simplest model to explain 5’ intron bias is homologous recombination between a reverse transcript of an intron-lacking mRNA and the original DNA locus to erase introns in the coding region (63). A similar mechanism could simultaneously erase IESs in coding regions via germline recombination between the MIC chromosome and a reverse transcript; however, they are usually in different subcellular locations. More plausibly, a source for DNA recombination could be a MAC nanochromosome, already abundant at high copy number, but another source could be a reverse transcript of a long non-coding template RNA that guides DNA rearrangement (14, 15). Either recombination event in the germline would lead to loss of IESs, while retaining introns, but neither would necessarily provide a bias for IES-loss in coding regions. The 5’ bias of IESs could also reflect an evolutionary bias for continuous coding regions. Alternatively, upstream IESs might regulate gene expression or cell growth (50, 64, 65).

We identified at least 174 examples of overlapping intron and IES positions across the three species, supporting an IES-intron conversion model proposed in (33). This suggests a potential fluidity between intron- and IES-removal in ciliates, two noncoding regions removed at different stages. Ciliate genomes are generally intron-sparse. *Oxytricha* averages 1.7 introns/gene, *Tetmemena* has 1.1 and *E. woodruffi* has 2.2. The fact that *E. woodruffi* has the most introns but the smallest number of IESs per gene (Figure 3) suggests that many non-coding insertions are either removed at different stages or less abundant in the germline, as is the case for 3’ IESs in *E. woodruffi*. The intron-sparseness of ciliates is compatible with a hypothesis that, given the opportunity, it is advantageous to eliminate non-coding sequences earlier at the DNA level. The overlap between hundreds of introns and IESs is consistent with a possible error correction opportunity to remove DNA insertions as introns if they fail to be excised as IESs, either during development or evolution (33).

This study investigated the evolution of scrambled genes by comparing *Oxytricha* and *Tetmemena* to *E. woodruffi*, as an earlier-diverged representative of the spirotrich lineage. While *E. woodruffi* has approximately half as many scrambled genes as *Tetmemena* and *Oxytricha*, its genes are also much more continuous. For example, the most scrambled gene in *E. woodruffi,* encoding a DNA replication licensing factor (EUPWOO_MAC_28518, 3kb), has only 20 scrambled junctions. The most scrambled gene in *Tetmemena* (LASU02015934.1, 14.7kb, encoding a hydrocephalus-inducing-like protein) has 204 scrambled pointers, and the most scrambled gene in *Oxytricha* (Contig17454.0, 13.7kb, encoding a dynein heavy chain family protein, ref. 1) is similarly complex, with 195 scrambled junctions. Together, these observations are consistent with our interpretation that *E. woodruffi* reflects an evolutionary intermediate stage, as it contains both fewer scrambled loci and fewer scrambled junctions within its scrambled loci. The observation that the most scrambled locus differs in each species is also consistent with the conclusion that complex gene architectures may continue to elaborate independently.

We observed that scrambled genes in each species tend to have more paralogs than nonscrambled genes. Similarly, in *Chilodonella uncinata* (18), a distantly-related ciliate in the class *Phyllopharyngea* which also has scrambled genes, scrambled gene families (orthogroups) contain more genes (∼2.9) than nonscrambled gene families (∼1.3) (18). scrambled patterns could readily arise from local duplications (Figure S5). These observations are most consistent with a simple model (42) in which local duplications permit combinatorial DNA rearrangement between paralogous germline regions, and mutation accumulation in either paralog establishes a scrambled pattern that can propagate by weaving together segments from paralogous sources. Other proposed models include Hoffman and Prescott’s (66) IES-invasion model that suggested that pairs of IESs could invade an MDS, and then subsequently recombine with another IES to yield odd/even scrambled regions; however, a previous examination did not find support for this model (33). Prescott *et al.* (67) also proposed that some odd-even scrambled loci could arise suddenly via reciprocal recombination with loops of AT-rich DNA. We proposed a gradual model in which MDS/IES recombination at short AT-rich repeats (precursors to pointers) could generate and propagate odd-even scrambled patterns (33, 55). While limited comparisons of orthologs favored the stepwise recombination models (32, 33, 34, 35), none of the earlier models accounted for the widespread occurrence of partial paralogy, revealed by genome assemblies.

## Methods

### DNA collection and sequencing of *Tetmemena sp*

*Tetmemena sp.* (strain SeJ-2015; ref. 7) was isolated as a single cell from a stock culture and propagated as a clonal strain via vegetative (asexual) cell culture. Cells were cultured in Pringsheim media (0.11mM Na_2_HPO_4_, 0.08mM MgSO_4_, 0.85mM Ca(NO_3_)_2_, 0.35mM KCl, pH 7.0) and fed with *Chlamydomonas reinhardtii*, together with 0.1%(v/v) of an overnight culture of non-virulent *Klebsiella pneumoniae*. Macronuclei and micronuclei were isolated using sucrose gradient centrifugation (68). Genomic DNA was subsequently purified using the Nucleospin Tissue Kit (Takara Bio USA, Inc.). Macronuclear DNA was sequenced and assembled in Chen *et al.* (7). Micronuclear DNA was further size-selected via BluePippin (Sage Science) for PacBio sequencing, or via 0.6% (w/v) SeaKem Gold agarose electrophoresis (Lonza) for Illumina sequencing. Micronuclear DNA purification and sequencing protocols are described in (1).

### DNA collection and sequencing for *E. woodruffi*

*E. woodruffi* (strain Iz01) was cultured in Volvic water at room temperature and fed with green algae every 2-3 days. We fed cells with *Chlamydomonas reinhardtii* for MAC DNA collection, and switched to *Chlorogonium capillatum* for MIC DNA collection. In order to remove algal contamination, cells were starved for at least 2-3 days before collection. Cells were washed and concentrated as in Chen et al. (1). Because MAC DNA is predominant in whole cell DNA, we used whole cell DNA (purified via NucleoSpin Tissue kit, Takara Bio USA, Inc.) for MAC genome sequencing. Paired-end sequencing was performed on an Illumina Hiseq2000 at the Princeton University Genomics Core Facility.

MIC DNA was enriched from whole cell DNA and sequenced via three sequencing platforms (Illumina, Pacific Biosciences and Oxford Nanopore Technologies). We used conventional and pulse-field gel electrophoresis to enrich MIC DNA:

1. High-molecular-weight DNA was separated from whole cell DNA by gel-electrophoresis (0.25% agarose gel at 4 °C, 120 V for 4 hr). The top band was cut from the gel and purified with the QIAGEN QIAquick kit. The purified high-molecular-weight DNA was directly sent to the group of Dr. Robert Sebra at the Icahn School of Medicine at Mount Sinai for library construction and sequencing. BluePippin (Sage Science) separation was used before sequencing to select DNA >10kb. DNA was sequenced on two platforms: Illumina HiSeq2500 (150 bp paired-end reads) and PacBio Sequel (SMRT reads).
2. High-molecular-weight DNA was also enriched by pulsed-field gel electrophoresis (PFGE). *E. woodruffi* cells were mixed with 1% low-melt agarose to form plugs according to Akematsu et al. (69), with addition of 1 hr incubation with 50 μg/ml RNase (Invitrogen AM2288) in 10 mM Tris-HCl (pH7.5) at 37 °C for RNA depletion. After three washes of 1 hr with 1x TE buffer, the DNA plugs were incubated in 1mM PMSF to inactivate proteinase K, followed by MspJI (New England Biolabs) digestion at ^m^CNNR(9/13) sites to remove contaminant DNA (^m^C indicates C5-methylation or C5-hydroxymethylation). Previous reports have shown that no methylcytosine is detectable in vegetative cells of *Oxytricha* (70), *Tetrahymena* (71) and *Paramecium* (72), suggesting that C5-methylation and C5-hydroxymethylation are rarely involved in the vegetative growth of the ciliate lineage. We also validated by qPCR that the quantity of two randomly selected MIC loci is not changed after the MspJI digestion. On the contrary, algal genomic DNA is significantly digested by MspJI. Based on these results, we conclude that MspJI digestion can be used to remove bacterial and algal DNA with C5-methylation and C5-hydroxymethylation, leaving *E. woodruffi* MIC DNA intact. The agarose plugs containing digested DNA were then inserted into wells of 1.0% Certified Megabase agarose gel (Bio-Rad) for PFGE (CHEF-DR II System, Bio-Rad). The DNA was separated at 6 V, 14°C with 0.5X TBE buffer at a 120° angle for 24 hr with switch time of 60-120 seconds. We validated by qPCR that the *E. woodruffi* MIC chromosomes were not mobilized from the well, while the MAC DNA migrated into the gel. The MIC DNA was then extracted by phenol-chloroform purification. Library preparation and sequencing were performed at Oxford Nanopore Technologies (New York, NY).

### MAC genome assembly of *E. woodruffi*

We assembled the MAC genome of *E.woodruffi* using the same pipeline for *Tetmemena sp.* (7) for comparative analysis: two draft genomes were assembled by SPAdes (73) and Trinity (74), and were then merged by CAP3 (75). Trinity, which is software developed for *de novo* transcriptome assembly (74), has been used to assemble hypotrich MAC genomes (7) because their nanochromosome genome structure is similar to transcriptomes, including properties such as variable copy number and alternative isoforms (10). Telomeric reads were mapped to contigs by BLAT (76), and contigs were further extended and capped by telomeres when at least 5 reads pile up at a position near ends by custom python scripts (https://github.com/yifeng-evo/Oxytricha_Tetmemena_Euplotes/tree/main/MAC_genome_telomere_capping). The mitochondrial DNA was removed if the contig has a TBLASTX (77) hit on the *Oxytricha* mitochondrial genome (Genbank accession JN383842.1 and JN383843.1) or two *Euplotes* mitochondrial genomes (*Euplotes minuta* GQ903130.1, *Euplotes crassus* GQ903131.1). Algal contigs were removed by BLASTN to all *Chlamydomonas reinhardtii* nucleotide sequences downloaded from Genbank. Non-telomeric contigs were mapped to bacterial NR by BLASTX to remove bacterial contaminations. The genome was further compressed by CD-HIT (78) in two steps: 1) contigs <500bp were removed if 90% of the short contig can be aligned to a contig >=500bp with 90% similarity (-c 0.9 -aS 0.9 -uS 0.1); 2) Then the genome was compressed by 95% similarity (-c 0.95 -aS 0.9 -uS 0.1). Contigs shorter than 500bp without telomeres were removed. Nine contigs, likely Tec contaminants from the MIC genome, were also excluded (Tblastn, “-db_gencode 10 -evalue 1e-5”), and they could be assembled due to the high copy number in the MIC genome (46, 47, Genbank accessions of Tec ORFs are AAA62601.1, AAA62602.1, AAA62603.1, AAA91339.1, AAA91340.1, AAA91341.1, AAA91342.1).

### RNA sequencing of *E. woodruffi*

Total RNA was isolated from asexually growing *E. woodruffi* cells using TRIzol reagent (Thermo Fisher Scientific), and enriched for the poly(A)+ fraction using the NEBNext® Poly(A) mRNA Magnetic Isolation Module (New England Biolabs). Stranded RNA-seq libraries were constructed using the ScriptSeq v2 RNA-seq library preparation kit (Epicentre) and sequenced on an Illumina Nextseq500 at the Columbia Genome Center. The transcriptome was assembled by Trinity (74) and transcript alignments to the MAC genome were generated by PASA (79).

### Gene prediction of the *E. woodruffi* MAC genome and validation of MAC genome completeness

We followed the gene prediction pipeline developed by the Broad institute (https://github.com/PASApipeline/PASApipeline/wiki) using EVidenceModeler (EVM, 80) to generate the final gene predictions. EVM produced gene structures by weighted combination of evidence from three resources: *ab initio* prediction, protein alignments and transcript alignments (the weight was 3, 3, 10 respectively). *Ab initio* prediction was generated by BRAKER2 pipeline (81). Protein alignments for EVM were generated by mapping *Oxytricha* proteins to the *E. woodruffi* MAC genome by Exonerate (82). EVM predicted 33379 genes on MAC chromosomes with at least one telomere.

We assessed MAC genome completeness using three methods: 1) 28294 (80.6%) of the 35099 *E. woodruffi* MAC contigs have at least one telomere. 2) In the *E. woodruffi* genes predicted on telomeric contigs, 76.3% of BUSCO (83) eukaryotic genes were identified as complete. Within the 303 BUSCO genes, 205 are complete and single-copy, 26 are complete and duplicated, 20 are fragmented and 52 are missing. 3) We identified 51 tRNA encoding all 20 amino acids by tRNAscan-SE (84) in the MAC genome, including two suppressor tRNAs of UAA and UAG.

### MIC genome assembly of *Tetmemena sp*

The MIC genome of *Tetmemena* was assembled with a hybrid approach to combine reads from different sequencing platforms. *Tetmemena* Illumina reads were first assembled by SPAdes (73, parameters “-k 21,33,55,77,99,127 –careful”). PacBio reads were error corrected by FMLRC (85) using Illumina reads with default parameters. Corrected PacBio reads were aligned to both the MAC genome and the Illumina MIC assembly with BLASTN. Reads were removed if they start or end with telomeres or are aligned better to the MAC. The remaining reads were assembled with wtdbg2 (86, parameters “-x rs”). The PacBio assembly was polished by Pilon (87) with the “--diploid” option. The Illumina and PacBio assemblies were merged by quickmerge (88) with the “-l 5000” option.

### MIC genome assembly of *E. woodruffi*

The MIC genome of *E. woodruffi* was assembled using a similar procedure as described above for *Tetmemena*. *E. woodruffi* reads were filtered to remove bacterial contamination, including abundant high-GC contaminants, possibly endosymbionts (89). Nanopore reads with GC content >= 55% were assembled by Flye (90) with the parameter “--meta” for metagenomic assembly of bacterial contigs. We used kaiju (91) to identify bacteria taxa for these contigs. 9 of 10 top-covered contigs derive from Proteobacteria, from which many *Euplotes* symbionts derive (89). Bacterial contamination was removed from Illumina reads if perfectly mapping to these metagenomic contigs by Bowtie2 (92). The cleaned Illumina reads were then assembled by SPAdes with “-k 21,33,55,77,99,127” (73). Pacbio raw reads and Nanopore raw reads with GC content < 55% were aligned to a concatenated database containing both the MAC genome and the Illumina MIC assembly with BLASTN. Reads were removed if they start or end with telomeres or align better to the MAC. Remaining PacBio/Nanopore reads were assembled by Flye with “--meta” mode. The PacBio-Nanopore assembly was polished by Pilon with the “--diploid” option. Illumina and PacBio-Nanopore assemblies were merged by quickmerge with the “-l 10000” option. Contigs shorter than 1kb were removed.

### MIC genome decontamination

The draft MIC genome of *Tetmemena* was first mapped to telomeric MAC contigs by BLASTN. MIC contigs containing MDSs were included in the final assembly. The rest of the MIC contigs were filtered by a decontamination pipeline: 1) contigs were aligned to the *Klebsiella pneumoniae* genome, *Chlamydomonas reinhardtii* genome and the *Oxytricha* mitochondrial genome by BLASTN to remove contaminants; 2) the remaining contigs were then searched against the bacteria NR database and a ciliate protein database (including protein sequences annotated in *Tetrahymena thermophila*: http://www.ciliate.org/system/downloads/tet-latest/4-Protein%20fasta.fasta; *Paramecium tetraurelia*: http://paramecium.cgm.cnrs-gif.fr; and *Oxytricha trifallax*: https://oxy.ciliate.org) by BLASTX. Contigs with higher bit score to bacteria NR or G+C >45% were removed. The *E. woodruffi* MIC genome was decontaminated similarly, with addition of all *Chlorogonium* sequences (the algal food source) on NCBI and the two *Euplotes* mitochondrial genomes (*Euplotes minuta* GQ903130.1, *Euplotes crassus* GQ903131.1) to filter contaminants.

### Repeat identification

The repeat content in the MIC genomes was identified by RepeatModeler 1.0.10 (93) and RepeatMasker 4.0.7 (94) with default parameters.

### TBE/Tec detection

Representative *Oxytricha* TBE ORFs (Genbank accession AAB42034.1, AAB42016.1 and AAB42018.1) were used as queries to search TBEs in the *Oxytricha* and *Tetmemena* MIC genomes by TBLASTN (-db_gencode 6 -evalue 1e-7 -max_target_seqs 30000). Tec ORFs were similarly detected by using *Euplotes crassus* Tec1 and Tec2 ORFs as queries (-db_gencode 10 - evalue 1e-5 -max_target_seqs 30000, Genbank accessions of Tec ORFs are AAA62601.1, AAA62602.1, AAA62603.1, AAA91339.1, AAA91340.1, AAA91341.1, AAA91342.1). Complete TBEs/Tecs were determined by custom python scripts when three ORFs are within 2000 bp from each other and in correct orientation (https://github.com/yifeng-evo/Oxytricha_Tetmemena_Euplotes/tree/main/TBE_ORFs/TBE_to_oxy_genome_tblastn_parse.py, 43). 30 TBE ORFs with >70% completeness were subsampled from each species for phylogenetic analysis (except for the 57kD ORF in *Tetmemena*, for which 21 were subsampled). The subsampled TBE ORFs were aligned using MUSCLE (95) and the alignments were trimmed by trimAl “-automated1” (96). Phylogenetic trees were constructed using PhyML 3.3 (97).

### Rearrangement annotations

SDRAP (52) was used to annotate MDSs, pointers and MIC-specific regions (minimum percent identity for preliminary match annotation=95, minimum percent identity for additional match annotation=90, minimum length of pointer annotation=2). SDRAP requires MAC and MIC genomes as input. For the SDRAP annotation of *Oxytricha*, we used the MAC genome from Swart et al. (6) instead of the latest hybrid assembly that incorporated PacBio reads (10), because the former version was primarily based on Illumina reads, similar to the MAC genomes of *Tetmemena* (7, Genbank GCA_001273295.2) and *E. woodruffi* which are also Illumina assemblies. *Oxytricha* and *Tetmemena* MAC genomes were preprocessed by removing MAC contigs with TBE ORFs, considered MIC contaminants (43). SDRAP is a new program that can output the rearrangement annotations with minor differences from Chen et al. (1) but most annotations are robust (Figure S4). Scrambled and nonscrambled junctions/IESs were annotated by custom python scripts (https://github.com/yifeng-evo/Oxytricha_Tetmemena_Euplotes/tree/main/scrambled_nonscrambled_IES_pointer).

### MIC genome categories

Each MIC genome region is assigned to only one category in Figure 2A-C, even if it belongs to more than one category. The assignment is based on the following priority: MDS, TBE/Tec, MIC genes (only available for *Oxytricha*, which has developmental RNA-seq data), IES, tandem repeats, other repeats and non-coding non-repetitive regions. For example, a MIC region can be a TBE in an IES, and it is only considered as TBE in Figure 2A-C.

### Ortholog comparison pipeline and Monte Carlo simulations

Orthogroups of genes on telomeric MAC contigs were detected by OrthoFinder with “-S blast” (98). Single-copy orthologs were aligned by Clustal Omega (99). Protein alignments were reversely translated to CDS alignments by a modified script of pal2nal (100, https://github.com/yifeng-evo/Oxytricha_Tetmemena_Euplotes/tree/main/Ortholog_comparison/pal2nal.pl). Two modifications were made in the script: 1) the modified script allows pal2nal to take different genetic codes for three sequences (-codontable 6,6,10); 2) the script also fixed an error in the original pal2nal script in which codontable 10 for the Euplotid nuclear code was the same as the universal code. Visualization of pointer positions and intron locations on orthologs was implemented by a custom python script (https://github.com/yifeng-evo/Oxytricha_Tetmemena_Euplotes/blob/main/Ortholog_comparison/visualization_of_ortholog_comparison.py). Pointer positions or intron locations are considered conserved if they are within a 20bp alignment window on the CDS alignment. Protein domains were annotated by HMMER (101). We performed Monte Carlo simulations by randomly shuffling pointer locations on the CDS but keeping their original position distribution. This was implemented by a custom python script, which transforms the CDS to a circle, rotates pointer positions on the circle and outputs the shuffled position on the re-linearized CDS (https://github.com/yifeng-evo/Oxytricha_Tetmemena_Euplotes/blob/main/Ortholog_comparison/shuffle_simulation.py). The null hypothesis of the Monte Carlo test is that pointers positions are conserved by chance. *P*-value of Monte Carlo test is given by N_expected>observed_/N_total_ (N_expected>observed_ is the number of simulations when there are more conserved pointers in the simulation than the observation from real data, N_total_ =1000 in this study).

### PCR validation of Russian doll locus

The complex Russian doll locus on MIC contig TMEMEN_MIC_21461 in *Tetmemena* was validated by PCR to confirm the *Tetmemena* MIC genome assembly. *Tetmemena* micronuclear DNA was purified as described previously and used as template for PCR using PrimeSTAR Max DNA polymerase (Takara Bio). 11 primer sets (Table S12) were designed to amplify products between 3 kb and 6 kb in length, with overlapping regions between consecutive primer pairs. The resulting PCR products were visualized through agarose gel electrophoresis and bands of the expected size were extracted using a Monarch® DNA Gel Extraction Kit (New England Biolabs). The purified gel bands were cloned using a TOPO™ XL-2 Complete PCR Cloning Kit (Invitrogen), transformed into One Shot™ OmniMAX™ 2 T1R *E. coli* cells (Invitrogen), and individual clones were grown and their plasmids harvested with a QIAprep Spin Miniprep Kit (QIAGEN). The plasmid ends were Sanger sequenced, as well as the region where the *Oxytricha* MIC assembly contains inserted MDSs (Genewiz). Sanger sequencing reads were mapped to the *Tetmemena* MIC contig TMEMEN_MIC_21461 and visualized using Geneious Prime® 2021.1.1 (https://www.geneious.com).

## Acknowledgements

We thank Toshinobu Suzaki (Kobe University) for the gift of *E. woodruffi* (strain Iz01 from Shizuoka Prefecture) and *Chlorogonium capillatum*. We thank Sheela George for laboratory support and help with cell collection. We thank David Dai, Eoghan Harrington, John Beaulaurier and Sissel Juul at Oxford Nanopore Technologies in New York for providing sequencing and advice. We thank Robert Sebra and Melissa Smith for advice on PacBio sequencing. We thank Takahiko Akematsu, Lorraine Symington and Lea Marie for helping with PFGE. We thank Kaiyi Zhu, Shaojie He, Molly Przeworski, Harmen Bussemaker and Nataša Jonoska for advice on Monte Carlo simulation. We also thank Samuel Sternberg, Bill Jack, Scott Roy, and all current and past Landweber lab members for discussion about the origin of scrambled genes and Margarita T. Angelova, Sindhuja Devanapally, Danylo Villano and Kehan Bao for comments on the manuscript. This work was supported by the National Institutes of Health, R35GM122555, and National Science Foundation, DMS1764366, and the National Center for Genome Analysis Support computing resources (supported by National Science Foundation DBI1062432, ABI1458641, and ABI1759906 to Indiana University). Rafik Neme was supported by the Pew Latin American Fellows Program.

## Author contributions

Conceptualization: Y.F., R.N., L.Y.B. and L.F.L.; Methodology: Y.F., R.N. and L.Y.B.; Software: Y.F., X.C. and J.B.; Investigation: Y.F., L.Y.B and M.L.; Writing -Original Draft: Y.F.; Writing - Review & Editing: Y.F. and L.F.L.; Funding Acquisition: L.F.L.; Supervision: L.F.L.

