## Supplemental tables and figures for "Comparative genomics reveals insight into the evolutionary origin of massively scrambled genomes"

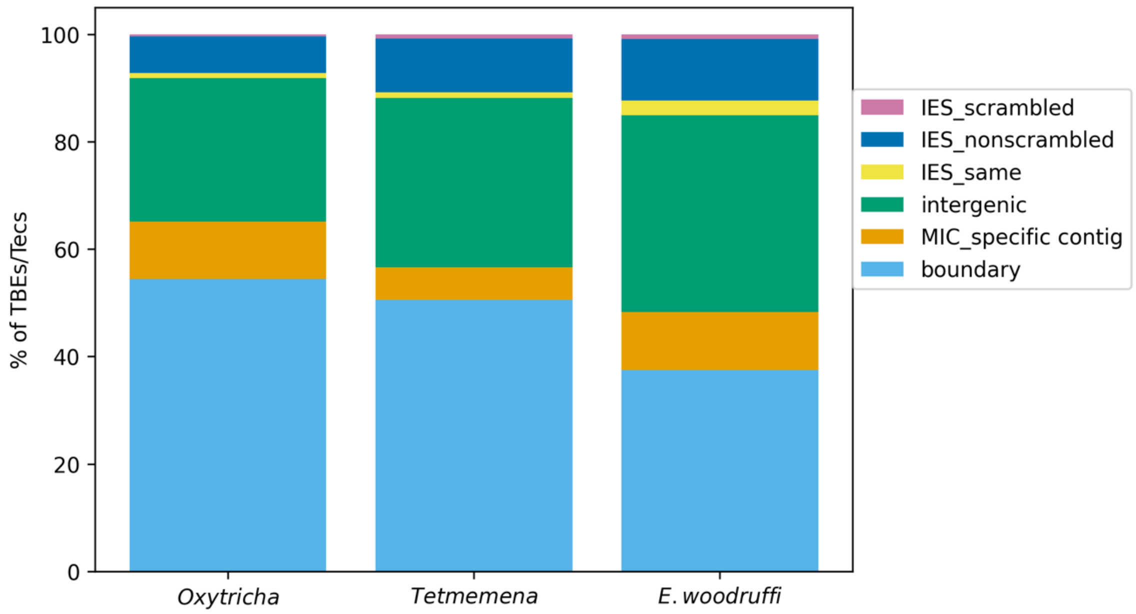


**Figure S1.** Comparison of MIC genome context of TBE/Tec transposons in three species. Complete and partial TBEs/Tecs were annotated by MIC regions. Boundary: boundaries of assembled MIC contigs. MIC-specific contig: no MDS identified on the MIC contig. Intergenic: MIC regions between MDSs for different MAC contigs. IES same: TE insertions between duplicated MDSs. IES nonscrambled: TE insertions between consecutive nonscrambled MDSs for the same MAC contigs. IES scrambled: MIC regions between nonconsecutive (scrambled) MDSs for the same MAC contig.

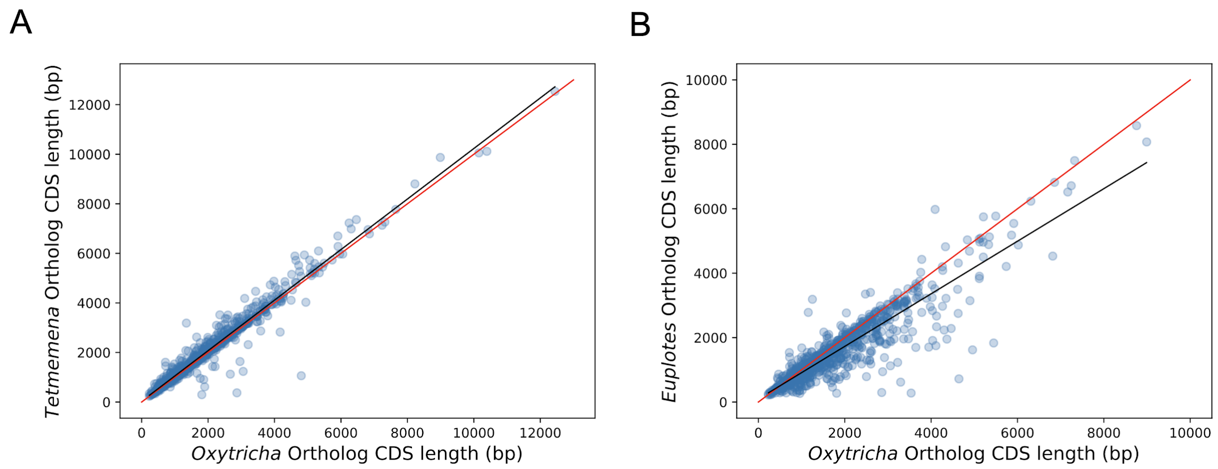


**Figure S2.** CDS lengths correlate for *Oxytricha*, *Tetmemena* and *E. woodruffi* orthologs (related to Figure 3). A) *Tetmemena* CDS length positively correlates with that of *Oxytricha* orthologs ($R^2$=0.96). Black line is the linear regression fitting function. Red line shows y=x. B) *E. woodruffi* CDS length positively correlates with that of *Oxytricha* orthologs ($R^2$=0.83).

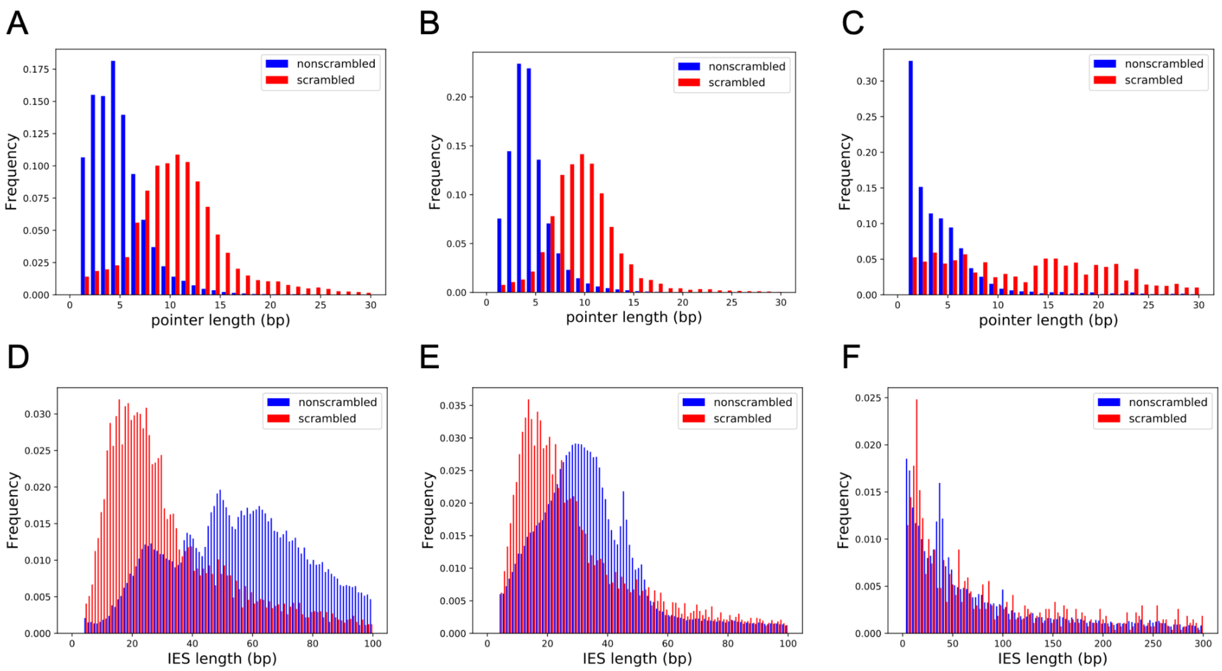


**Figure S3.** Scrambled and nonscrambled loci have distinct length distributions of IESs and pointers. A-C) Length distribution of scrambled and nonscrambled pointers <= 30bp in A) *Oxytricha*, B) *Tetmemena* and C) *E. woodruffi*. D- F) Length distribution of scrambled and nonscrambled IESs in D) *Oxytricha* (<=100bp), E) *Tetmemena* (<=100bp) and F) *E. woodruffi* (<=300bp).

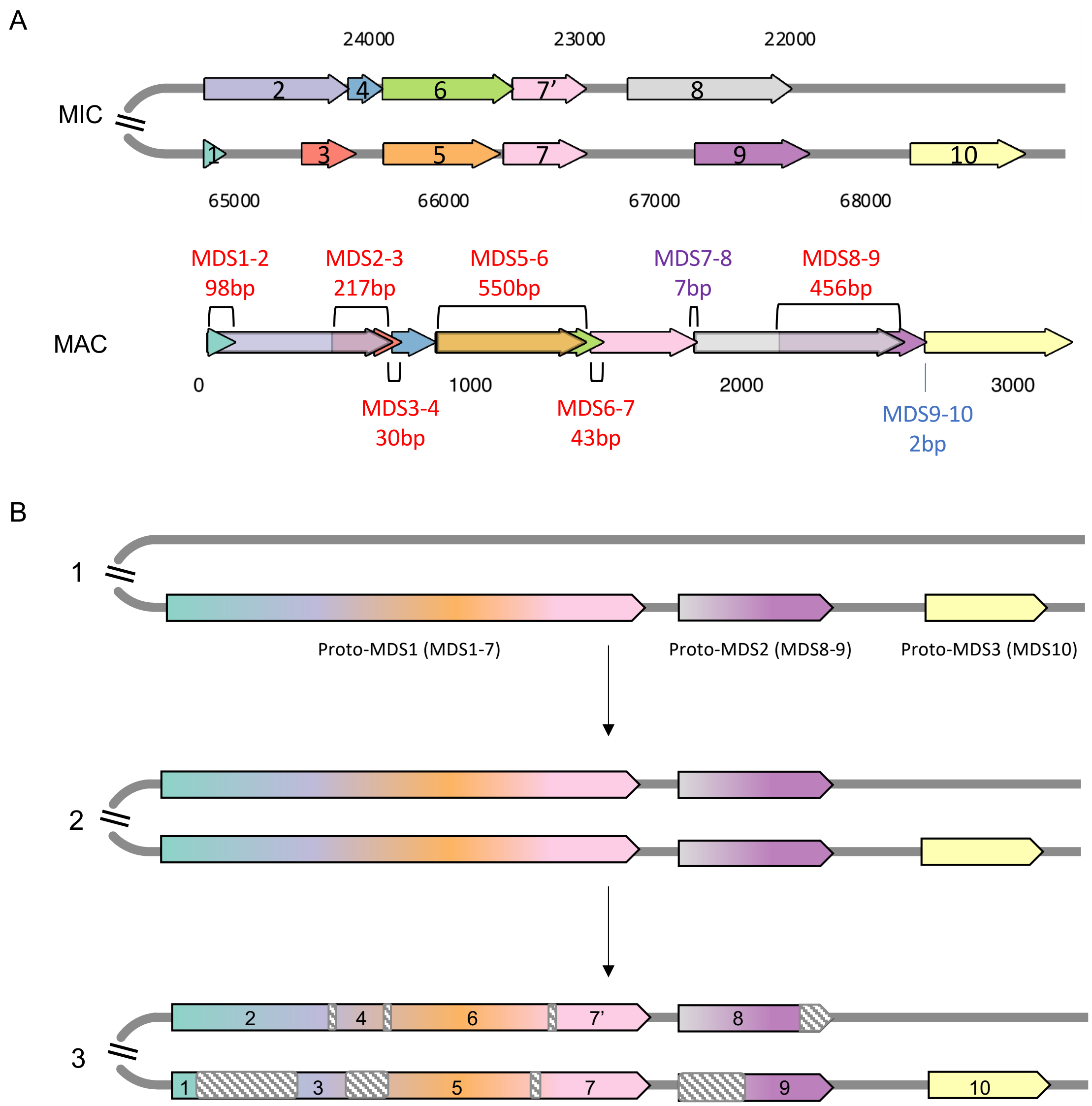


**Figure S4.** An example of an *E. woodruffi* scrambled gene locus containing paralogous MDSs. A) The upper panel is the map of a scrambled MIC locus (EUPWOO_MIC_17325). Below is the corresponding map of the MAC chromosome (EUPWOO_MAC_29939). Pointers between MDSs are labeled above or below the MAC contig (nonscrambled pointer length in blue and scrambled pointers labeled in red). B) A model for the evolutionary origin of this scrambled MIC locus by partial duplication and decay (adapted from [1]).

Stage 1: The ancestral MIC locus contains three nonscrambled MDSs (labeled proto-MDSs because they are precursors for the modern state). Stage 2: The region containing two proto-MDSs duplicated in the MIC genome. Stage 3: Nucleotide substitutions accumulated in both paralogous copies at different positions (shown in gray dashed boxes) leading to the fixation of some regions as MDSs, while the regions that accumulated more mutations decayed into IESs, which are removed during genome rearrangement.

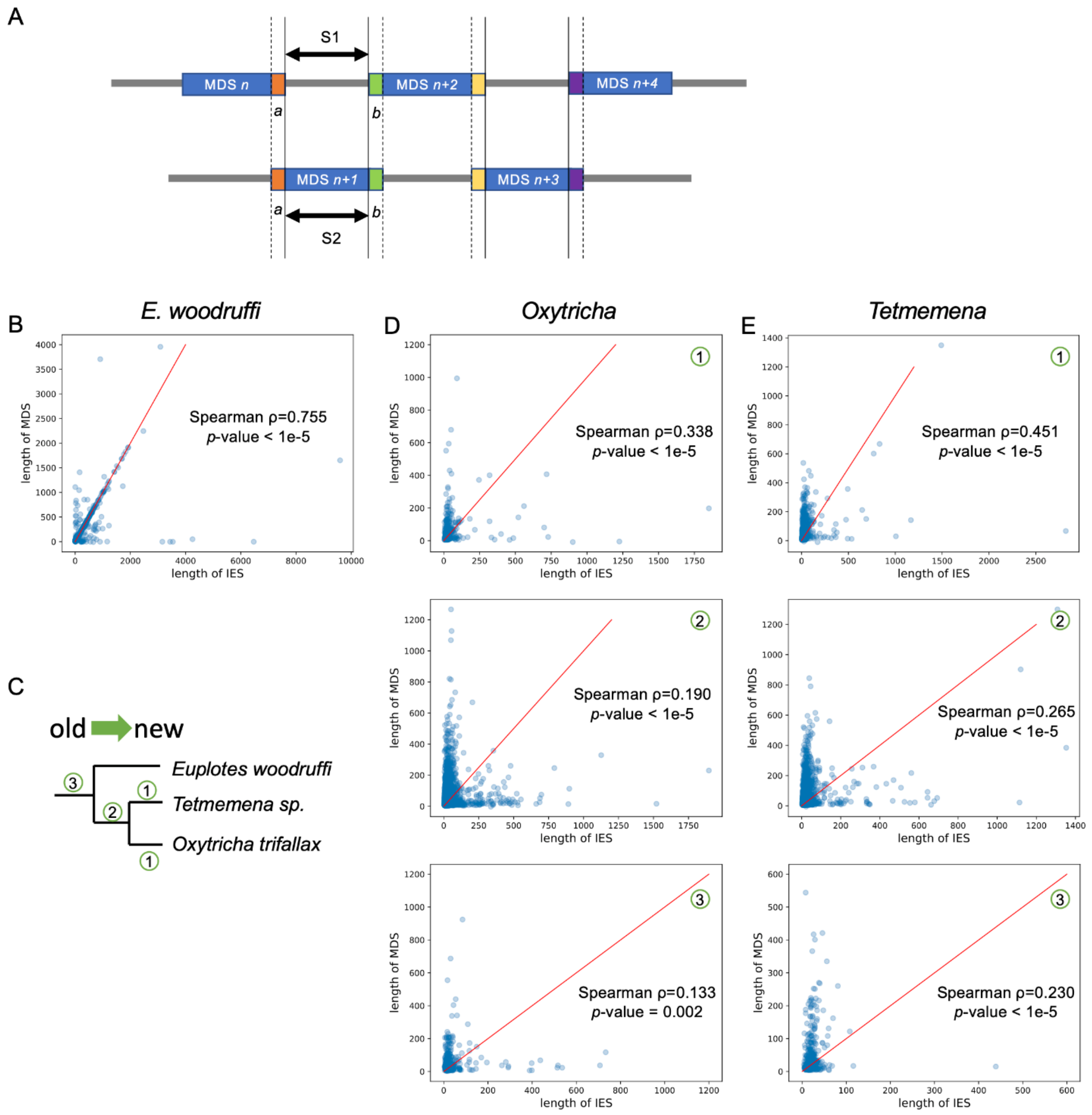


**Figure S5.** The trend of scrambled loci to contain odd-even patterns may arise from partial duplication followed by mutation accumulation. A) A diagram describing a typical scrambled region with an odd-even pattern. We propose that the IES (S1) between MDS *n* and MDS *n+2* may be ancestrally paralogous to MDS *n+1* (S2) which evolved by duplication of MDS *n+1* before it was scrambled. S1 and S2 would therefore be homologous in this model. B) The lengths of modern IES (S1) and MDS (S2) display a strong positive correlation in *E. woodruffi* (504 pairs). Many data points fall on the y=x (red line). All MDS and IES pairs were only considered if they are on the same MIC contig, to exclude alleles. C) Character mapping of scrambled loci on a phylogeny: 1. Examples of scrambled loci uniquely present in one species (only showing for *Oxytricha* and *Tetmemena*; most scrambled genes in *E. woodruffi* have no ortholog detectable in the other two species, possibly because the long genetic distance obscured homology, see main text and Table S3); 2. Scrambled loci shared between *Oxytricha* and *Tetmemena*, but not *E. woodruffi*; 3. Scrambled loci shared in three species. The lengths of IES (S1) and MDS (S2) in typical odd-even regions display a moderately positive correlation in *Oxytricha* (D) and *Tetmemena* (E). Newer scrambled loci correlate more strongly. Red line represents y=x. Note that S1 and S2 are flanked by identical pointers, *a* and *b*, in all annotated pairs.

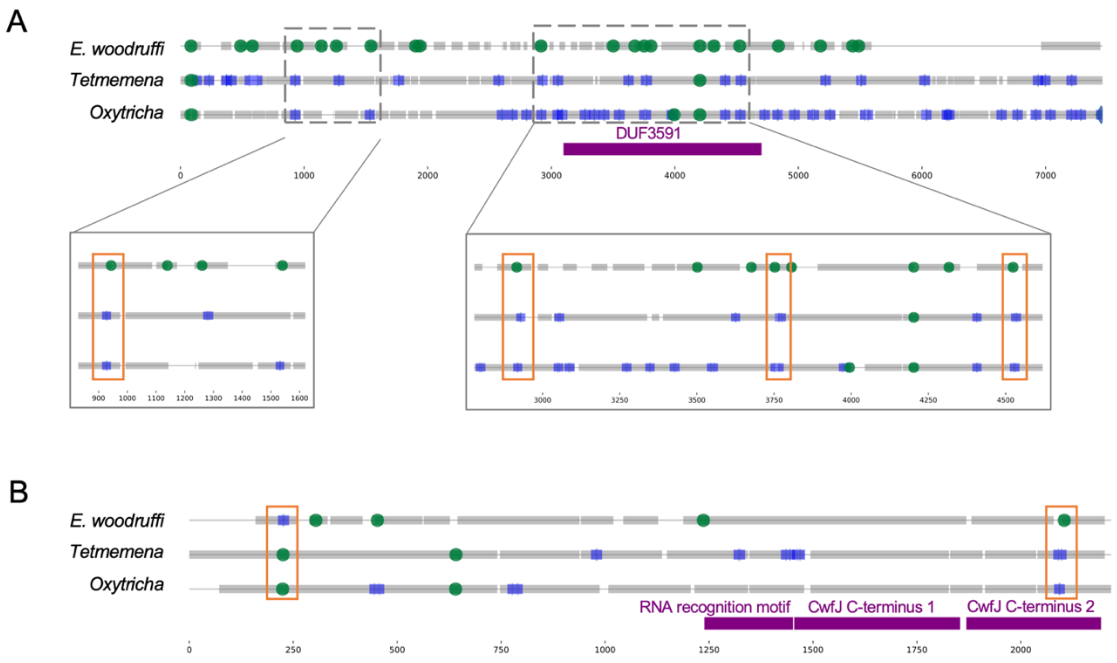


**Figure S6.** Examples of Intron-IES conversion across three species. A) Four intron positions in *E. woodruffi* (orange boxes in magnified regions) overlap locations of nonscrambled pointers in the orthologous genes in *Oxytricha* and *Tetmemena* (*Oxytricha*: Contig13378.0.g40; *Tetmemena*: LASU02004100.1.g1; *E. woodruffi*: EUPWOO_MAC_08218.g1) consistent with a possible trend of some ancestral introns becoming IESs in the hypotrich lineage. Two positions fall within a conserved protein domain of unknown function (DUF3591). B) An orthologous gene with two intron-IES conversions in reciprocal directions (*Oxytricha*: Contig16930.0.g77; *Tetmemena*: LASU02013377.1.g1; *E. woodruffi*: EUPWOO_MAC_15089.g1). Colors and annotation as in Figure 5.

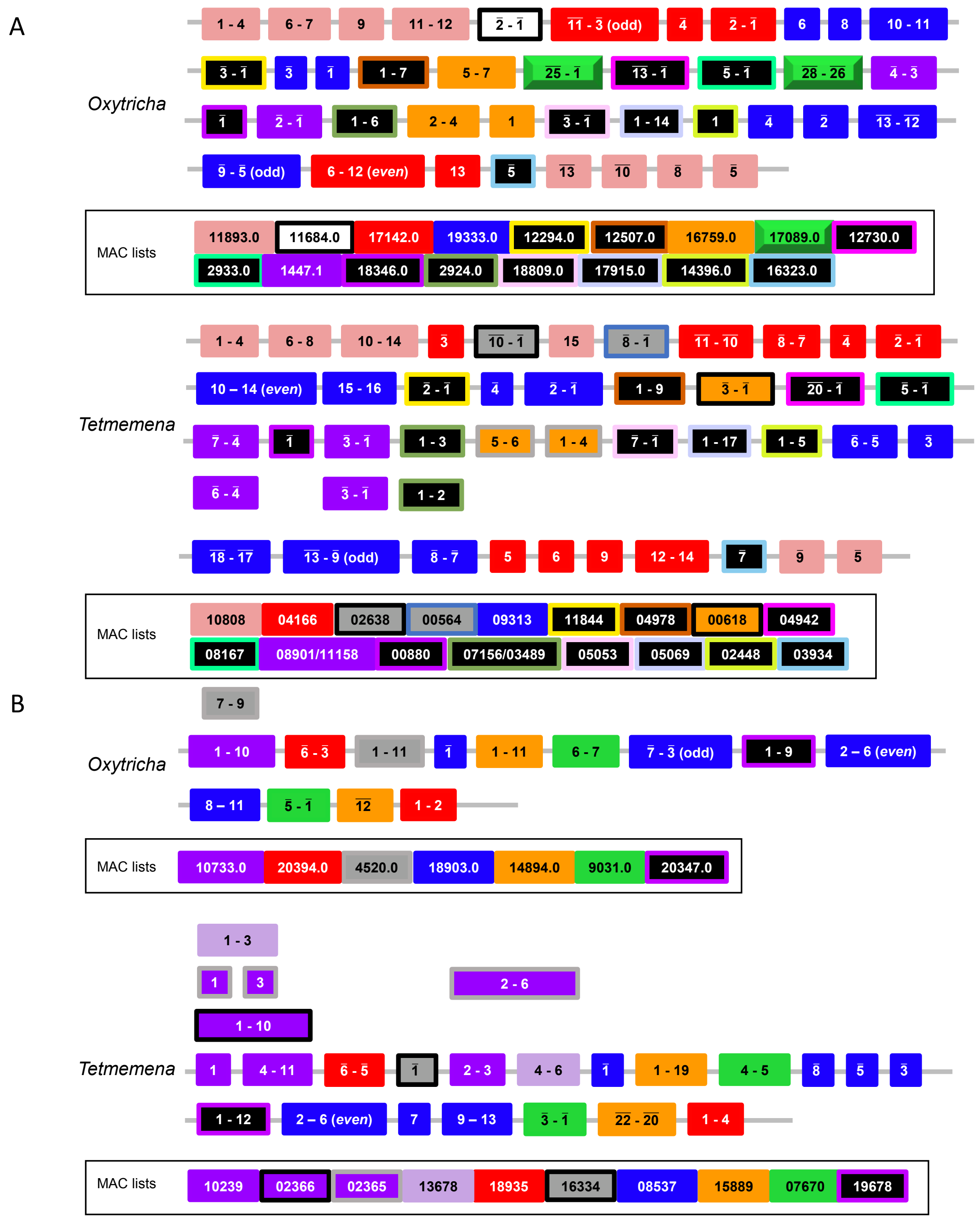

**Figure S7.** Detailed illustration of both Russian Doll regions in Figure 6. MDS indices are annotated here for each MAC locus. Overlined numbers represent inverted MDSs. MAC contig numbers for the MDSs are listed below and shown in corresponding color patterns (the *Oxytricha* loci were previously characterized in [2]).

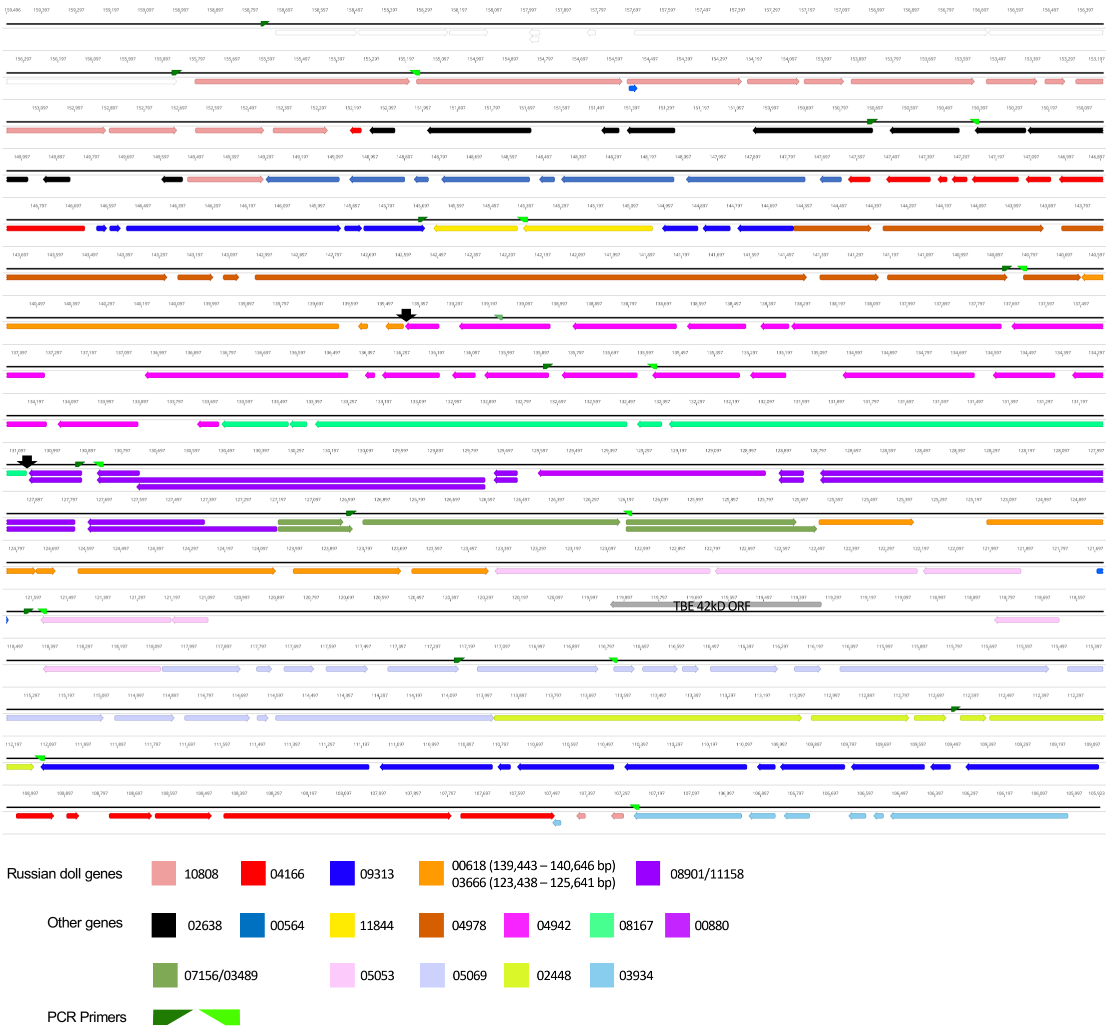


**Figure S8.** Details of the Russian Doll region in *Tetmemena* (TMEMEN_MIC_21461, Figure 6A). The whole region (~50 kb) was validated by 11 PCRs. The two black arrows indicate the absence of a Russian doll gene (green in Figure 6A) that is present in *Oxytricha*. Legend lists the 20 *Tetmemena* MAC contigs that contain the corresponding MDSs.

**Table S1.** Sequencing depth statistics for MIC genome assemblies

| | ***Oxytricha trifallax**** | ***Tetmemena sp.*** | ***Euplotes woodruffi*** |
|---|---|---|---|
| Illumina coverage (X) | 118 | 69 | 190 |
| PacBio coverage (X) | 34 | 28 | 24 |
| Nanopore coverage (X) | - | - | 41 |

*Sequencing data from Chen et al. [3].

**Raw reads were mapped to the MIC genome assembly by Minimap2 [4] and Bowtie2 [5]. Average coverage was calculated with BBmap pileup.sh [6] for MDS-containing contigs in the MIC genome assembly.

**Table S2.** Subcategories of repeat content in the three species.

| | | ***Oxytricha trifallax*** | ***Tetmemena sp.*** | ***Euplotes woodruffi*** |
|---|---|---|---|---|
| Transposable elements | Class 2 TBE/Tec | 15.4% | 1.9% | 2.3% |
| | Class 2 cut-and-paste DNA transposons (excluding TBE/Tec) | 2.8% | 0.9% | 2.8% |
| | Class 1 LTR | 1.2% | 1.1% | 0.7% |
| | Class 1 LINE | 1.1% | 0.1% | 0.2% |
| | Class 2 *Helitron* | 0.5% | 0.0% | 0.1% |
| | Class 1 SINE | 0.1% | 0.0% | 0.0% |
| | Unclassified | 27.4% | 14.9% | 15.9% |
| Tandem repeats | Satellites | 0.1% | 0.1% | 0.0% |
| | Simple repeats | 1.4% | 1.0% | 0.5% |
| | Low-complexity | 0.4% | 0.4% | 0.2% |
| Total percentage of the MIC genome assembly | | 49.6% | 20.2% | 22.5% |

Repeat content of the three genomes, as annotated by Repeatmasker [7] with additional manual annotation of TBE/Tec elements. The numbers may differ from Figure 2A-C because some repeats are assigned as other MIC categories in the pie charts (Methods). For example, a MIC region which is both an IES and satellite, is assigned as IES in Figure 2A-C, but is counted as a satellite in this table.

**Table S3.** TBE/Tec ORFs in three species

| | ***Oxytricha trifallax*** | ***Tetmemena sp.*** | ***Euplotes woodruffi*** |
|---|---|---|---|
| ORF1 | 21,433 | 1645 | 1202 |
| ORF2 | 16,816 | 723 | 112 |
| ORF3 | 19,582 | 641 | 245 |
| complete | 9313* | 48 | 74 |

* Differs from 10,109 in Chen et al. [3] because we used different versions of BLAST and custom python scripts to identify complete TBEs (See Methods).

**Table S4.** Orthology among scrambled and nonscrambled genes in the three species

| | *Oxytricha trifallax* | | | *Tetmemena sp.* | | | *Euplotes woodruffi* | | | ciliate database* | |
|---|---|---|---|---|---|---|---|---|---|---|---|
| | scrambled | non-scrambled | No ortholog | scrambled | non-scrambled | No ortholog | scrambled | nons-crambled | No ortholog | yes | no |
| *Oxytricha* scrambled genes (3613) | | | | 51.26% | 28.62% | 20.12% | 3.43% | 23.89% | 72.68% | 82.12% | 17.88% |
| *Tetmemena* scrambled genes (3371) | 50.25% | 21.77% | 27.97% | | | | 3.14% | 21.89% | 74.96% | 73.57% | 26.43% |
| *E. woodruffi* scrambled genes (2429) | 4.90% | 11.98% | 83.12% | 4.45% | 12.27% | 83.29% | | | | 68.67% | 31.33% |
| *Oxytricha* nonscrambled genes (19454) | | | | 5.88% | 72.43% | 21.69% | 2.22% | 21.74% | 76.04% | 78.91% | 21.09% |
| *Tetmemena* nonscrambled genes (21377) | 6.36% | 60.39% | 33.26% | | | | 1.81% | 18.74% | 79.46% | 67.44% | 32.56% |
| *E. woodruffi* nonscrambled genes (30950) | 3.44% | 13.58% | 82.98% | 3.17% | 14.25% | 82.58% | | | | 68.80% | 31.20% |

* Ciliate database is generated by extracting all protein sequences in phylum Ciliophora (taxid: 5878) from NR database.

**Table S5.** More scrambled MAC contigs contain at least one paralogous MDS that may be involved in alternative rearrangement.

| | ***O. trifallax*** | ***Tetmemena sp.*** | ***E.woodruffi*** |
|---|---|---|---|
| Paralogous MDSs on scrambled contigs | 694 | 894 | 441 |
| Paralogous MDSs on nonscrambled contigs | 1340 | 949 | 524 |
| Scrambled contigs with paralogous MDSs (vs. total scrambled contigs) | 304 (2852) | 270 (2556) | 223 (1913) |
| Nonscrambled contigs with paralogous MDSs (vs. total nonscrambled contigs) | 809 (17484) | 435 (18609) | 422 (26381) |
| *p*-value (chi-square test) for contig numbers | 4e-39 | 1e-104 | 4e-177 |

**Table S6.** MDS-IES pairs share homologous sequences in the three species (related to Figure S5).

| | *E. woodruffi* | | *Oxytricha* | | | *Tetmemena* | | |
|---|---|---|---|---|---|---|---|---|
| | | | recent -> ancestral | | | recent -> ancestral | | |
| | similar length[a] | all | Group 1[b] | Group 2 | Group 3 | Group 1 | Group 2 | Group 3 |
| # of MDS-IES pairs | 248 | 504 | 370 | 4055 | 520 | 1005 | 3222 | 369 |
| # of pairs with a homologous core sequence | 224 | 272 | 17 | 20 | 2 | 16 | 20 | 2 |
| Ratio | 90.32% | 53.97% | 4.59% | 0.49% | 0.38% | 1.59% | 0.62% | 0.54% |
| G test with Williams correction *p-value* | - | - | 9.09e-09 | | | 0.022 | | |

[a] MDS length is between 0.8 * IES length and 1.2 * IES length.

[b] MDS-IES pairs were dated on a phylogenetic tree as shown in Figure S5C.

**Table S7.** Presence of conserved pointers in three species, with Monte Carlo simulations

| | | *Oxytricha* vs. *Tetmemena* | *Oxytricha* vs. *Euplotes* | *Tetmemena* vs. *Euplotes* |
|---|---|---|---|---|
| Observed # of conserved pointers | Number of orthologs | 1345 | 51 | 52 |
| | Number of pointer pairs | 4448 | 56 | 58 |
| | Pointers conserved in 3 species | 23 | | |
| Expectations (Monte Carlo simulations) | Number of orthologs | 697 | 52 | 57 |
| | *p*-value | <0.001 | 0.59 | 0.738 |
| | Number of pointer pairs | 1781 | 57 | 62 |
| | *p*-value | <0.001 | 0.6 | 0.783 |
| | Pointers conserved in 3 species | 6.7 | | |
| | *p*-value | <0.001 | | |

**Table S8.** Scrambled pointers are more conserved than nonscrambled pointers.

| | *Oxytricha* | | *Tetmemena* | |
|---|---|---|---|---|
| | scrambled | nonscrambled | scrambled | nonscrambled |
| Conserved | 1442 | 2715 | 1412 | 2549 |
| Unconserved | 611 | 6091 | 801 | 8046 |
| chi-square test *p*-value | 1e-239 | | 5e-296 | |

**Table S9.** Most pointers conserved in position are different in sequence

| | *Oxytricha-Tetmemena* pointers |
|---|---|
| Same | 389 |
| pointer 1 contained in pointer 2 | 320 |
| pointer 2 contained in pointer 1 | 250 |
| Different | 3489 |
| Total | 4448 |

**Table S10.** Intron-IES conversion comparison in three species and Monte Carlo simulations

| | *Euplotes* intron – *Oxytricha*/*Tetmemena* IES | | | *Tetmemena* intron – *Oxytricha*/*Euplotes* IES | | | *Oxytricha* intron – *Tetmemena*/*Euplotes* IES | | |
|---|---|---|---|---|---|---|---|---|---|
| | expected | observed | *p*-value | expected | observed | *p*-value | expected | observed | *p*-value |
| Positions | 34 | 103 | <0.001 | 1.4 | 1 | 0.767 | 1.8 | 0 | - |
| | *Euplotes* IES – *Oxytricha*/*Tetmemena* intron | | | *Tetmemena* IES – *Oxytricha*/*Euplotes* intron | | | *Oxytricha* IES – *Tetmemena*/*Euplotes* intron | | |
| | expected | observed | *p*-value | expected | observed | *p*-value | expected | observed | *p*-value |
| Positions | 2.4 | 24 | <0.001 | 11 | 34 | <0.001 | 7.6 | 12 | 0.098 |

**Table S11.** Pairwise intron-IES conversion comparisons and Monte Carlo simulations

| | ***Oxytricha* intron–*Tetmemena* IES** | | | ***Oxytricha* intron – *Euplotes* IES** | | | ***Tetmemena* intron– *Euplotes* IES** | | |
|---|---|---|---|---|---|---|---|---|---|
| | expected | observed | *p*-value | expected | observed | *p*-value | expected | observed | *p*-value |
| Positions | 416 | 463 | 0.008 | 29 | 31 | 0.402 | 26 | 26 | 0.55 |
| | ***Tetmemena* intron– *Oxytricha* IES** | | | ***Euplotes* intron – *Oxytricha* IES** | | | ***Euplotes* intron – *Tetmemena* IES** | | |
| | expected | observed | *p*-value | expected | observed | *p*-value | expected | observed | *p*-value |
| Positions | 308 | 247 | 1 | 288 | 277 | 0.745 | 349 | 347 | 0.559 |

**Table S12.** PCR primers for validation of the Russian doll region in *Tetmemena* MIC DNA (Figure 6A)

| Primer | Sequence |
|---|---|
| 1 F | CATTCTTATTTCCCTTCATTTGTTTC |
| 1 R | CTTTCAATCTATTAAGGAGTATCTC |
| 2 F | GGCTAAAGTAAGAATATTTTATTTGAAG |
| 2 R | CAATAAATGCATGAGTTTAAATAATATCG |
| 3 F | GAGCAGGCTTGATTCAACAAAATC |
| 3 R | CATTTAAATCTTAAAAAGAGATTTTTCC |
| 4 F | CACCTACTAACTTTGAAAGACAAAG |
| 4 R | CATAGAGCTGATTTAATAACTTCATATC |
| 5 F | CTGCCCAGTCCAAATTTAAATCAAT |
| 5 R | GTTGTTAATATTTCCTTACTTATTAC |
| 6 F | TATAGCAGCTAAGGAAATCAAAATTAG |
| 6 R | CTTTTAAAGAAGGGGACAAATAACAAG |
| 7 F | CTTACCAAAAGCATTATTTAAGATGC |
| 7 R | GGATCTAATAGTGTAATAAATATCTTG |
| 8 F | ACTTACACTCAATTTAAAACAGATTG |
| 8 R | CAGATTTTCCTCCATGTTTAAAAGTC |
| 9 F | GTTCACTATGAATCTAGAAGAGATTTAAG |
| 9 R | CTCTTTCCTGATTATTCAAGGAAAAATAG |
| 10 F | CATAAATCAGACTAAAAAATTCATGC |
| 10 R | CAAAATAGATATGATAATGTCAGAAATG |
| 11 F | CTTATGTCTCTAGTAAAAATAATTATAAAC |
| 11 R | GGCATTTCATAGATCTTACTTTAAC |
| 40 F | CTCGGTATACATATATAACTATG |

**Dataset S1 (separate file).** Pointers conserved in all three species.

**Dataset S2 (separate file).** The TBE pointers in *Oxytricha* that are conserved with non-TBE pointers in *Tetmemena.*
